# SOX4 Reprograms Adipose Stromal Cells into a Cancer-Associated Fibroblast- like State to Drive Metabolic Disease

**DOI:** 10.64898/2026.09.24.754142

**Authors:** Karima Drareni, Ryan P. Calhoun, Lan Cheng, Camille E. M. Piguet, Lin Liu, Sarah Traynor, Josephine M. Tan, Marco Angelozzi, David M. Merrick, Veronique Lefebvre, Patrick Seale

**Affiliations:** Institute for Diabetes, Obesity & Metabolism Perelman School of Medicine, University of Pennsylvania, Philadelphia, PA, USA, 19104; Department of Cell and Developmental Biology; Perelman School of Medicine, University of Pennsylvania, Philadelphia, PA, USA, 19104; Department of Medicine; Perelman School of Medicine, University of Pennsylvania, Philadelphia, PA, USA, 19104; Division of Orthopedics, Department of Surgery, Children’s Hospital of Philadelphia, Philadelphia, PA, USA

**Keywords:** Adipose Tissue, Obesity, Metabolic disease, cancer-associated fibroblast, mesenchymal stromal cell, SOX4, Midkine

## Abstract

Pathogenic adipose tissue remodeling promotes metabolic disease in obesity, but the mechanisms that establish this unhealthy tissue state remain poorly understood. Here, we show that obesity drives SOX4-dependent reprogramming of mesenchymal stromal cells (MSCs) into cancer-associated fibroblast-like (CAF-like) cells that promote adipose tissue dysfunction. TGFβ signaling is elevated in obesity and activates SOX4 in mouse and human MSCs, inducing their conversion to a CAF-like state. In mice, MSC-specific SOX4 activation induces the CAF-like program and exacerbates adipose tissue inflammation and glucose intolerance, whereas Sox4 deletion attenuates inflammation and improves glucose homeostasis during obesity. We further identify the growth factor Midkine (MDK) as a SOX4-regulated paracrine effector produced by CAF-like cells. MDK inhibition in obese mice reduces adipose tissue inflammation and improves metabolic function. Together, these findings define a TGFβ-SOX4-MDK stromal signaling axis that drives pathological adipose tissue remodeling in obesity and highlight this pathway as a potential therapeutic target for improving metabolic health.

## Introduction

Obesity increases the risk of numerous comorbidities, including type 2 diabetes, cardiovascular disease, stroke, and several cancers^1,2^. Many of these complications are closely linked to adipose tissue pathology and dysfunction^3–5^. Maladaptive adipose tissue remodeling in obesity is characterized by adipocyte hypertrophy, impaired adipogenesis, inflammation, and fibrosis. These pathological features compromise adipose tissue function and predispose to ectopic lipid deposition in other tissues and metabolic disease.^6,7^.

Adipose tissue growth and maintenance is governed by interactions among multiple cell types, including adipocytes, immune cells, vascular endothelial cells, and mesenchymal stromal cells (MSCs)^3,7,8^. Among these, MSCs play many important roles in regulating adipose tissue function by providing structural support, producing extracellular matrix components, and serving as adipocyte progenitors. MSCs also dynamically regulate immune responses in adipose and other tissues^9–11^. However, under chronic metabolic stress such as obesity, MSCs may acquire maladaptive inflammatory and fibrotic states that contribute to adipose tissue dysfunction^10,12^. Recent research has drawn attention to parallels between the adipose tissue microenvironment in obesity and the tumor microenvironment in cancer^13,14^. In tumors, stromal remodeling driven by cancer-associated fibroblasts (CAFs) promotes fibrosis, immune modulation, and disease progression.

Here, we show that obesity induces SOX4-dependent CAF-like cells that drive metabolically unhealthy remodeling of adipose tissue. TGFβ signaling, which is increased in adipose tissue during obesity, activates SOX4 expression in DPP4⁺ progenitor MSCs, driving their transition toward a pro-inflammatory state while suppressing adipogenic differentiation. Experimental activation of SOX4 in MSCs promotes adipose tissue inflammation and exacerbates metabolic dysfunction. We further found that CAF-like cells secrete Midkine (MDK), a heparin-binding growth factor previously implicated in cancer, tissue regeneration, and immune modulation. MDK promotes inflammatory remodeling of adipose tissue and impairs systemic metabolism, whereas MDK inhibition ameliorates adipose tissue pathology and improves metabolic homeostasis. Together, these findings identify a SOX4-dependent pathway that drives pathogenic stromal reprogramming in adipose tissue and suggest a therapeutic avenue for improving metabolic health.

## Results

### High fat diet induces a cancer-associated fibroblast-like population in adipose tissue

To investigate stromal-mechanisms that regulate maladaptive adipose tissue remodeling during obesity, we performed single-cell RNA sequencing (scRNA-seq) on the stromal vascular fraction (SVF) isolated from epididymal white adipose tissue (eWAT) of C57BL/6 mice fed either a control diet (CD) or high-fat diet (HFD) for 16 weeks. eWAT is a visceral fat depot that undergoes pronounced pathological remodeling during obesity^3,5^. Unsupervised clustering resolved the major cell types, with cluster identities assigned using canonical lineage markers, including MSCs (*Pdgfra*), macrophages (*Adgre1*), T cells (*Cd3e*), B cells (*Cd79a*), dendritic cells (*Flt3*), endothelial cells (*Cldn5*), mast cells (*Cpa3*), and mesothelial cells (*Krt19*) (**Fig. 1a**; **Extended data Fig. 1a, b**). As expected, HFD feeding led to marked expansion and modulation of immune cell populations (**Fig. 1b**; **Extended data Fig. 1c**). In particular, the macrophage compartment shifted away from resident/homeostatic states toward a pro-inflammatory profile, with increased lipid-associated macrophages (LAMs) (**Extended data Fig. S1d**).

**Fig. 1.**
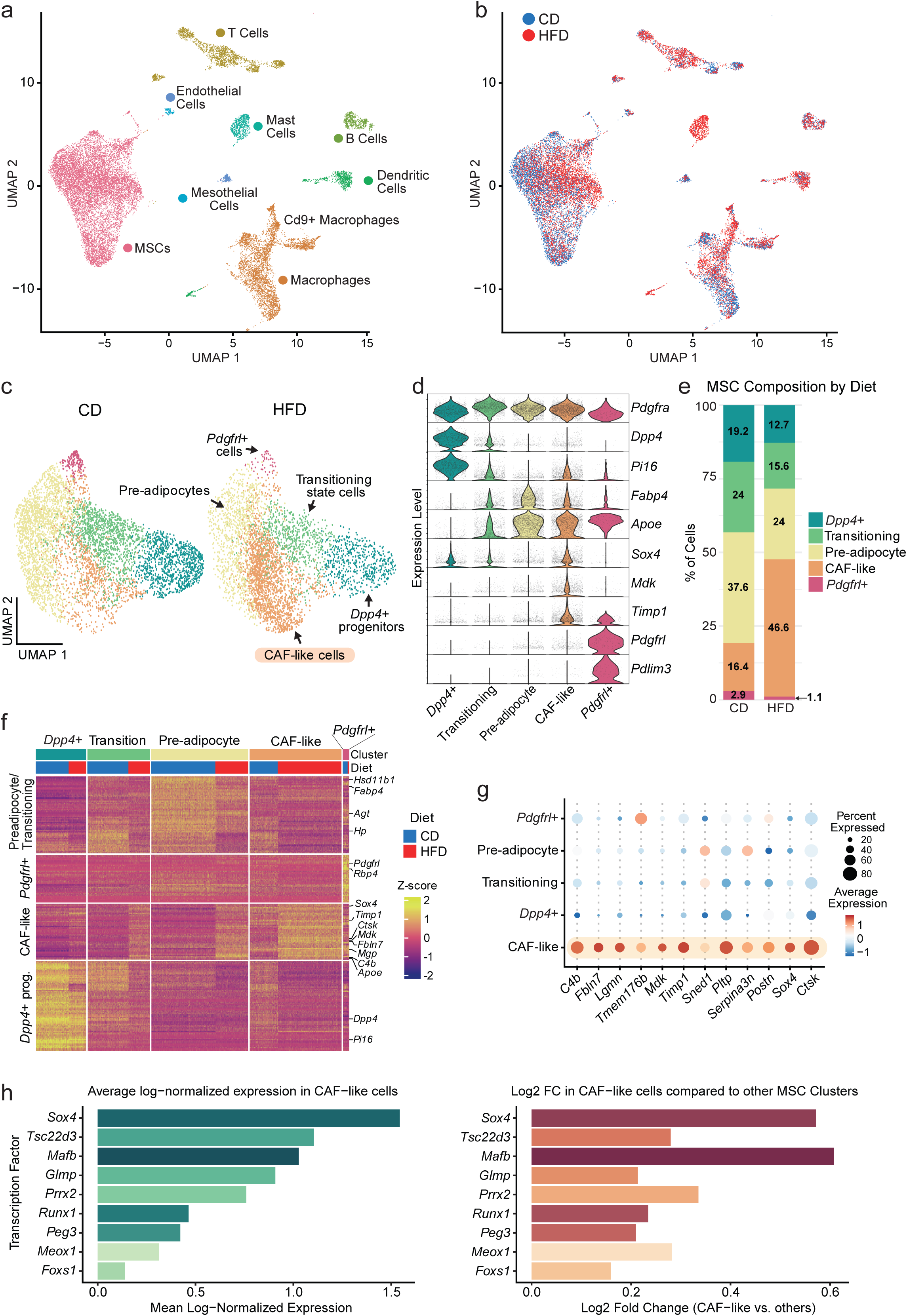
Obesity induces a CAF-like stromal cell population in epididymal white adipose tissue. **(a)** Integrated UMAP representation of 18,975 stromal vascular fraction (SVF) cells from eWAT of mice fed CD or HFD for 16 weeks, annotated by major cell types. **(b)** UMAP from (a), colored by diet condition (CD, n=9440, blue; HFD, n=9535, red). **(c)** UMAP showing MSCs (*Pdgfra*^+^) split by diet condition, identifying *Dpp4^+^* progenitors, transitioning cells, pre-adipocytes, CAF-like cells, and *Pdgfrl^+^*cells. **(d)** Violin plots showing expression levels of representative marker genes across MSC subsets. **(e)** Stacked bar plot showing the relative abundance of MSC subsets under CD and HFD conditions. **(f)** Expression heatmap of cluster-enriched genes across MSC subsets and diet conditions. **(g)** Dot plot showing expression of CAF-like cell marker genes across MSC subsets. **(h)** Transcription factors with enriched expression in CAF-like cells, shown as mean log-normalized expression (left panel) and log2 fold change relative to other MSC clusters (right panel). n = 3 mice per diet group pooled for scRNA-seq analysis.

We focused our analysis on MSCs by reclustering this compartment, identifying five transcriptionally distinct subpopulations: *Dpp4*⁺ progenitors, transitioning state cells, pre-adipocytes, *Pdgfrl*⁺ cells, and a CAF-like population (**Fig. 1c**). *Dpp4*^+^ progenitors were marked by expression of *Dpp4* and *Pi16* (**Fig. 1d, f**). Pre-adipocytes expressed enriched levels of adipogenic genes such as *Fabp4* and *Hsd11b1* (**Fig. 1d, f**). *Pdgfrl*^+^ cells expressed *Pdgfrl* and other fibroblast-associated genes such as *Rbp4* and *Pdlim3* (**Fig. 1d, f**). CAF-like cells were distinguished by their expression of many inflammatory and extracellular matrix-related genes, including *Mdk* and *Timp1* (**Fig. 1d, f, g**). Obesity strongly altered the composition of the MSC compartment. CAF-like cells were rare under CD conditions (∼16% of MSCs) but were greatly expanded in obesity (HFD conditions) (∼46% of MSCs) (**Fig. 1e**). This expansion of CAF-like cells was accompanied by a relative reduction in other populations, particularly transitioning cells and pre-adipocytes, suggesting that obesogenic conditions shift MSC states away from adipogenesis and toward a CAF-like phenotype (**Fig. 1e**).

Pathway analysis of the genes enriched in CAF-like cells identified processes associated with inflammation, matrix remodeling, migration and adhesion. Consistent with this profile, these cells expressed many canonical CAF-associated genes, including *C4b*, *Fbln7*, *Mdk*, *Timp1*, *Postn*, *Ctsk*, and *Sox4* (**Fig. 1f, g; Extended Data Fig. 1e**). We next compared CAF-like cells with previously described adipose stromal populations associated with fibrosis and inflammation. The fibro- inflammatory progenitor (FIP) signature described by Hepler *et al.*^10^ was predominantly enriched in DPP4^+^ MSCs, with limited expression in CAF-like cells, whereas the CD9-high fibroblast signature described by Marcelin *et al.* ^12^ was most strongly enriched in the *Pdgfrl*^+^ population, with a subset of CD9-high signature genes also expressed in CAF-like cells (**Extended Data Fig. 1f**). Thus, CAF-like cells are transcriptionally distinct from these previously described stromal populations.

To further assess how obesity alters MSC identity, we examined the expression of cluster-selective genes across CD and HFD conditions. Obesity reduced the expression of cluster-defining genes in pre-adipocytes and transitioning state cells while inducing CAF-like marker genes, including *Sox4*, *Timp1*, *Ctsk*, *Mdk*, and *Mgp*, across multiple MSC populations (**Fig. 1f**). We next examined transcription factor expression to identify candidate regulators of the CAF-like state. This analysis revealed *Sox4* as one of the most highly enriched and abundant transcription factors in CAF-like cells (**Fig. 1h**). Altogether, these results demonstrate that HFD-induced obesity shifts MSC fate away from an adipogenic trajectory and toward a CAF-like inflammatory state.

### SOX4 marks and drives CAF-like cell identity in adipose tissue during obesity

To determine the timing of SOX4 activation and CAF-like cell induction during obesity, we analyzed *Sox4* mRNA expression in eWAT over the course of HFD feeding. *Sox4* expression progressively increased during HFD feeding and was significantly elevated at 12 and 16 weeks (**Fig. 2a**). Other CAF signature genes, such as *Fbln7*, *Mgp*, *Mdk* and *Ctsk* exhibited a similar temporal pattern, supporting a link between *Sox4* and the CAF-like gene program induced by obesity (**Extended Data Fig. 2a**). *Sox4* and CAF-like gene expression were also increased in iWAT and rpWAT after 16 weeks of HFD feeding, with no changes observed in brown adipose tissue or liver (**Extended Data Fig. 2b, c**). Immunofluorescence staining for SOX4 and the pan-MSC marker PDGFRα identified few SOX4^+^ MSCs in the eWAT of CD-fed/lean mice. By contrast, obese mice (16 week HFD) exhibited a marked increase in SOX4⁺ cells distributed throughout the adipose tissue (**Fig. 2b**). Quantification revealed an approximate threefold increase in SOX4⁺ MSCs in obesity (**Fig. 2b**), consistent with the proportions predicted from scRNA-seq analysis. Taken together, these results indicate that obesity induces the progressive upregulation of SOX4 at both the mRNA and protein levels, accompanied by expansion of a CAF-like cell population in WAT.

**Fig. 2.**
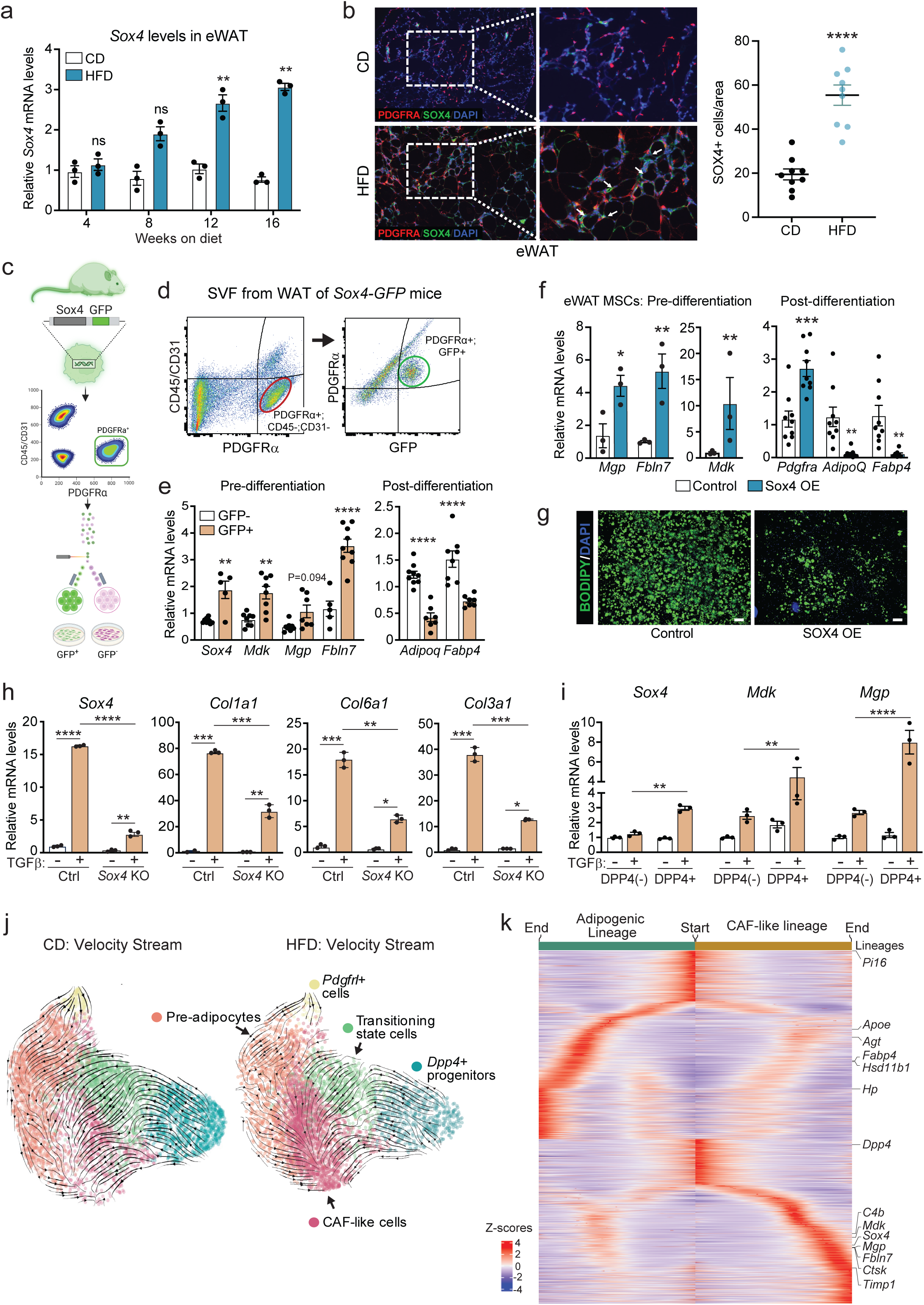
SOX4 is induced by obesity and promotes a CAF-like, non-adipogenic fate. **(a)** *Sox4* mRNA levels in eWAT after 4, 8, 12, and 16 weeks of HFD compared with C D feeding (n = 3 mice per group/time point). **(b)** Immunofluorescence staining of PDGFRα (red), SOX4 (green), and nuclei (DAPI, blue) in eWAT from CD- and HFD-fed mice. (Right) Quantification of SOX4+ cells normalized to total nuclei. (n = 3 mice per group). **(c)** Schematic of the *Sox4*-GFP transgene and strategy used to isolate *Sox4* (GFP)+ and *Sox4* (GFP)- MSCs. **(d)** Representative gating strategy for isolation of GFP+ and GFP- MSCs. **(e)** mRNA levels of: CAF-like genes in *Sox4* (GFP)+ and *Sox4* (GFP)- MSCs (Pre-differentiation, left), and adipocyte marker gene levels in *Sox4* (GFP)+ and *Sox4* (GFP)- cells after inducing adipogenic differentiation (right). (n = 6 per condition). **(f)** mRNA levels of: CAF-like genes in primary eWAT MSCs isolated from control and Sox4-expressing mice (Pre-differentiation, left) (n = 3 per group); and *Pdgfra*, *Adipoq*, and *Fabp4* in control and Sox4-expressing cells after inducing adipogenic differentiation (right) (n = 9 per condition). **(g)** Bodipy staining of triglycerides in control and SOX4-expressing primary eWAT MSCs following adipogenic differentiation. DAPI was used to stain nuclei (blue). **(h)** mRNA levels of *Sox4* and indicated collagen genes in control and *Sox4*- deficient 3T3-L1 cells following treatment with recombinant TGFβ or vehicle (n = 3 per condition). **(i)** mRNA levels of *Sox4*, *Mdk*, and *Mgp* in FACS-purified DPP4⁺ and DPP4⁻ MSCs isolated from eWAT of control mice and treated with recombinant TGFβ or vehicle (n = 3 per condition). **(j)** RNA velocity stream plots of eWAT MSCs from CD- and HFD-fed mice projected onto UMAP coordinates, showing predicted state transitions. Cells are colored by cluster identity. **(k)** Heatmap showing gene expression across pseudotime. Slingshot analysis identifies trajectories from *Dpp4*⁺ progenitors toward pre-adipocytes or CAF-like cells. Data are presented as mean ± SEM. Statistical significance: ns, not significant; *P < 0.05; **P < 0.01; ***P < 0.001; ****P < 0.0001.

Next, we utilized *Sox4-GFP* reporter mice to isolate SOX4^+^ MSCs. PDGFRα^+^; Sox4(GFP)^+^ and PDGFRα^+^; Sox4(GFP)^-^ cells were purified by fluorescence-activated cell sorting of stromal vascular cells from the eWAT of mice fed a HFD for 12 weeks (**Fig. 2c, d**). qRT-PCR analysis showed that Sox4(GFP)^+^ MSCs expressed higher levels of CAF-associated genes, including *Mdk*, *Mgp*, and *Fbln7* (**Fig. 2e**). To assess adipogenic capacity, sorted cells were treated with adipogenic differentiation inducers. Notably, Sox4(GFP)^+^ MSCs exhibited a significantly reduced capacity to undergo adipocyte differentiation compared with Sox4(GFP)^-^ MSCs, as evidenced by lower expression of adipocyte marker genes *Adipoq* and *Fabp4* (**Fig. 2e**).

To test whether SOX4 is sufficient to induce CAF-like cell identity and inhibit adipogenesis, we increased SOX4 expression in MSCs using *Pdgfra^CreERT2^*; *Rosa26^SOX4^* (SOX4-overexpressing [OE]) mice. In these mice, tamoxifen treatment activates Cre recombinase and induces SOX4 expression specifically in *Pdgfra*-expressing cells (i.e. MSCs). We isolated stromal vascular cells from the eWAT of SOX4-OE and control mice following tamoxifen treatment. SOX4 upregulation increased the levels of CAF-like cells signature genes *Mgp*, *Fbln7* and *Mdk*, under basal conditions (**Fig. 2f**). SOX4 expression also strongly inhibited adipogenic differentiation, as shown by markedly reduced lipid accumulation and an ∼80-90% decrease in the expression of adipocyte marker genes *Adipoq* and *Fabp4* in *Sox4*-expressing cells compared with control cells (**Fig. 2f, g**). SOX4- expressing cells also retained higher expression levels of the MSC marker *Pdgfra* under differentiation conditions (**Fig. 2f**). Consistent with these findings, lentiviral expression of SOX4 in 3T3-L1 pre-adipocytes induced CAF-like genes *Mdk*, *Mgp*, *Fbln7*, and impaired adipogenic differentiation (**Extended Data Fig. 2d-f**). Altogether, these findings show that SOX4 functions in MSCs to promote CAF-like cell fate and suppress adipogenic activity.

### TGFβ signaling activates SOX4 and the CAF-like program in DPP4^+^ progenitors

The cytokine TGFβ is elevated in adipose tissue during obesity and is an established repressor of adipogenesis and driver of fibrosis^15–19^. Interestingly, TGFβ has been shown to activate SOX4 expression in other contexts, particularly in cancer^20–23^. We found that TGFβ treatment of primary mouse MSCs robustly induced *Sox4* mRNA levels, whereas related SOXC family members *Sox11* and *Sox12* mRNA levels were unchanged (**Extended Data Fig. 2g**). TGFβ also upregulated canonical fibrosis-associated genes, including *Col1a1*, *Col3a1*, *Mmp9*, and *Mmp3* (**Extended Data Fig. 2g**). To determine whether SOX4 mediates TGFβ-induced fibrosis, we ablated *Sox4* in 3T3-L1 cells using CRISPR-Cas9, followed by TGFβ treatment. Notably, *Sox4*-deficiency markedly attenuated the induction of extracellular matrix and fibrotic genes, including *Col1a1*, *Col6a1*, and *Col3a1*, in response to TGFβ, but did not restore adipogenic gene induction under differentiation conditions (**Fig. 2h**; **Extended Data Fig. 2h**).

Prior work from our laboratory showed that DPP4^+^ progenitors are multipotent and more responsive to TGFβ signaling than other MSC populations^24,25^. This is consistent with broader evidence that MSC populations are organized along hierarchical lineage relationships across tissues, with DPP4^+^ cells designated as “universal progenitor fibroblasts”^26^. To determine whether TGFβ promotes CAF-like cell identity in DPP4^+^ cells, we purified DPP4^+^ and DPP4^-^ MSCs (all PDGFRα^+^) from eWAT and treated them with TGFβ. TGFβ activated CAF genes, such as *Sox4*, *Mdk*, and *Mgp*, to a much greater extent in DPP4^+^ cells relative to DPP4^-^ cells (**Fig. 2i**). Notably, *Sox4* was not induced by TGFβ in DPP4^-^ cells but was upregulated approximately threefold in DPP4+ cells (Fig. 2I). TGFβ upregulated *Mgp* by ∼8-fold in DPP4^+^ cells compared with ∼3-fold in DPP4^-^ cells (**Fig. 2j**). These results suggest that adipose tissue DPP4^+^ cells are susceptible to undergoing a phenotypic transition to a CAF-like state in response to elevated TGFβ signaling.

To determine whether DPP4^+^ progenitors may give rise to other MSC states during adipose tissue remodeling, we performed pseudotemporal trajectory analysis of the scRNA-seq data. This analysis predicted that DPP4^+^ MSCs progress through a transitional cell state before differentiating into either pre-adipocytes or CAF-like cells (**Fig. 2j**). Pre-adipocytes were also predicted to shift toward the CAF-like state during obesity (**Fig. 2j**). To visualize transcriptional changes along the inferred lineage trajectories, we generated a heatmap of genes regulated across pseudotime starting from DPP4^+^ progenitors (**Fig. 2k**). One branch showed progressive induction of adipogenic markers, including *Fabp4*, *Apoe*, and *Agt*, whereas the alternative branch displayed increased expression of *Sox4* and other CAF-related genes (e.g. *Mdk*, *Mgp*, *Timp1*, and *Fbln7*). Notably, cells on the adipogenic trajectory were predominantly derived from CD (lean) animals, whereas the CAF-like branch was enriched for cells from the HFD condition (**Fig. 2k**). These data suggest that DPP4^+^ cells give rise to either adipogenic cells or CAF-like cells, with obesity biasing lineage allocation toward the CAF-like fate.

### SOX4-activation in MSCs drives adipose tissue inflammation and glucose intolerance

We next examined the effects of MSC-selective SOX4 activation on adipose tissue remodeling and systemic metabolism using SOX4-OE mice. We validated transgene expression by immunoblot and qRT-PCR, which showed robust SOX4 induction in stromal-vascular cells isolated from eWAT and iWAT following tamoxifen treatment (**Extended Data Fig. 3a, b**). In addition, stromal vascular cells from both fat depots expressed higher levels of CAF-like genes, including *Mmp11*, *Mgp*, *Mdk*, *Fbln7*, *Ctsk*, and *Col1a1*, compared with controls (**Extended Data Fig. 3b**).

SOX4-OE and control mice were treated with tamoxifen to induce SOX4 expression in MSCs, maintained on CD for another 4 weeks, and then fed a HFD for 2 weeks (**Extended Data Fig. 3c**). Control and SOX4-OE mice showed comparable body weight trajectories throughout the study period (**Fig. 3a**). Weights of adipose tissue depots (eWAT, iWAT, rpWAT, and BAT) and liver were similar between Control and SOX4-OE mice at the final 6-week time point (**Fig. 3b**, **Extended Data Fig. 3d**). At 2- and 4- weeks of CD feeding, we did not detect significant changes in glucose tolerance, though there was a trend towards impairment in the SOX4-OE mice at 4 weeks (**Extended Data Fig. 3e**). Strikingly, following only 2 weeks of HFD exposure, SOX4-OE mice exhibited markedly impaired glucose tolerance compared to controls (**Fig. 3c**). Plasma insulin levels during the glucose tolerance test were comparable between control and SOX4-OE mice (**Fig. 3d**). Histological analysis revealed a more inflamed and fibrotic adipose tissue architecture in SOX4-OE mice (**Fig. 3e**). Adipocyte size in eWAT did not differ significantly between groups (**Fig. 3f**). Immunostaining for the macrophage marker Galectin-3 (Mac-2) revealed a ∼2.5-fold increase in the number of crown-like structures surrounding adipocytes in eWAT from SOX4-OE mice (**Fig. 3e, f**). CAF-associated genes *Mdk*, *Mgp*, *Fbln7*, *Col3a1*, and *Col6a1*, were upregulated by ∼4-12-fold in eWAT from SOX4- OE mice compared with controls (**Fig. 3g**). Consistent with the histological changes, eWAT from SOX4-OE mice exhibited increased expression of immune cell genes compared with controls, including ∼4-fold higher *Adgre1* (F4/80; macrophages) and ∼3-fold higher *Cd3e* (T cells) expression (**Fig. 3h**). *Adipoq* (adiponectin) expression was not significantly altered, whereas *Lep* (leptin) expression was increased in eWAT of SOX4-OE mice (**Fig. 3i**). iWAT from SOX4-OE mice showed increased expression of *Sox4* and *Mdk*, whereas other CAF-associated genes and immune markers were not significantly altered, and no overt histological changes were observed (**Extended Data Fig. 3f, g**). Livers from SOX4-OE mice expressed increased levels of inflammatory and fibrosis-related genes despite the absence of overt histological abnormalities (**Extended Data Fig. 3h, i**), suggesting that molecular alterations may precede detectable tissue pathology. We confirmed the effects of SOX4 activation in an independent cohort of SOX4-OE mice fed HFD for 2 weeks, showing similar effects of SOX4 in promoting inflammatory and CAF-like gene expression in eWAT, associated with and impaired glucose tolerance (**Extended Data Fig. 4**). In summary, SOX4 activation in MSCs induces inflammatory and fibrotic remodeling in eWAT and exacerbates metabolic dysfunction under obesogenic conditions without altering adiposity.

**Fig. 3.**
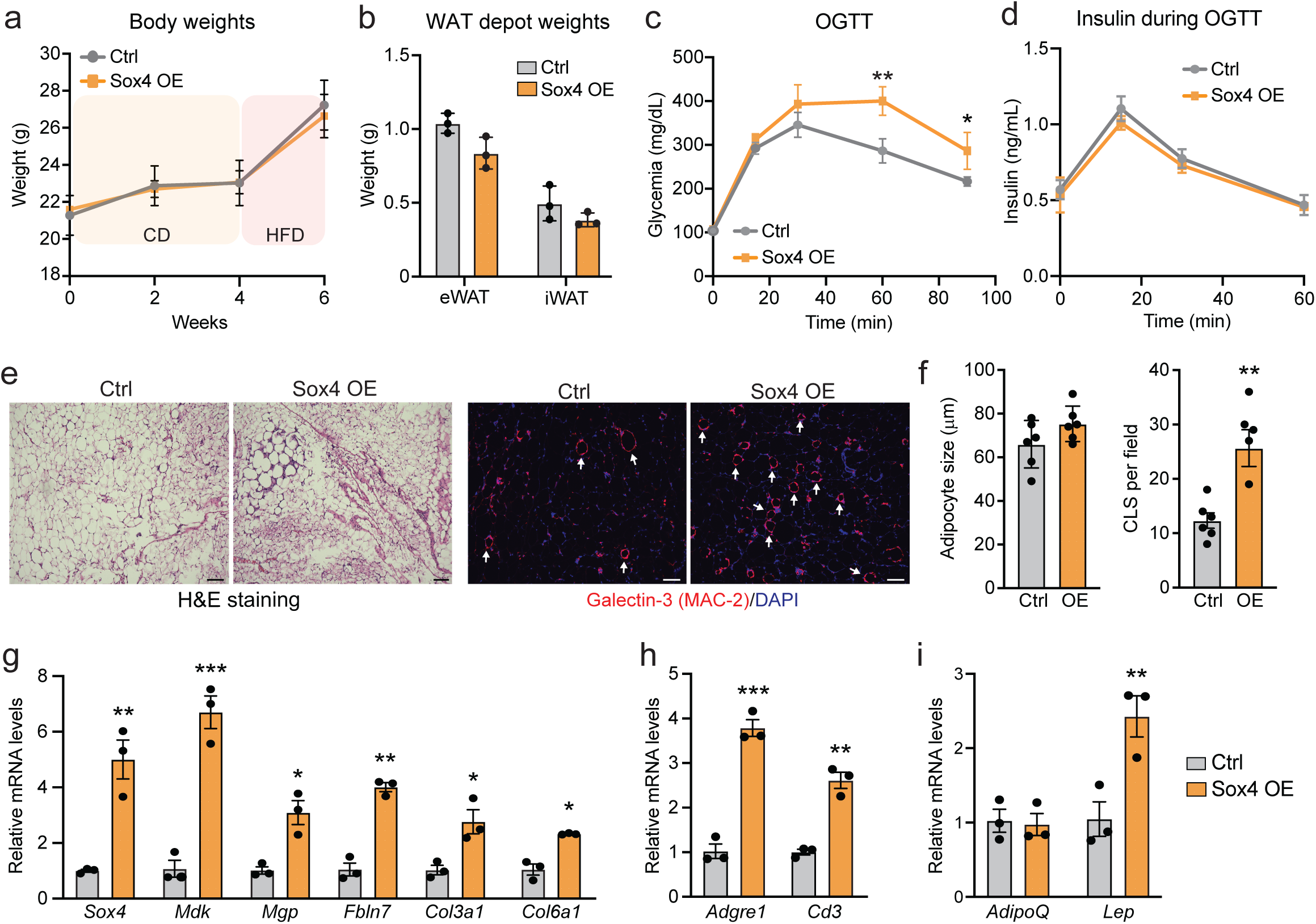
SOX4 activation in MSCs promotes inflammatory WAT remodeling and glucose intolerance. *Pdgfra^CreERT2^*; *Rosa26^Sox4^* (Sox4-OE) and control mice were treated with tamoxifen to activate SOX4 expression in MSCs, maintained on CD for 4 weeks and switched onto HFD for 2 weeks. **(a)** Body weight trajectories. **(b)** eWAT and iWAT depot weights. **(c)** OGTT. **(d)** Plasma insulin concentrations during OGTT. **(e)** H&E staining and immunofluorescence staining of Galectin-3 (Mac-2, red) and nuclei (DAPI, blue) in eWAT sections. **(f)** Quantification of adipocyte size and crown-like structures (CLS) in eWAT (n =6 fields per group from n = 3 mice per group). **(g)** mRNA levels of *Sox4* and CAF-like genes in eWAT. **(h)** mRNA levels of immune cell markers *Adgre1* and *Cd3* (n=3 mice per group). **(i)** mRNA levels of adipocyte-selective genes *Adipoq* and *Lep* in eWAT (n=3 mice per group). Data are presented as mean ± SEM. Statistical significance: *P < 0.05; **P < 0.01; ***P < 0.001.

### SOX4-deficiency in MSCs attenuates pathological adipose tissue remodeling and improves systemic metabolic function in obesity

We next tested if inhibition of SOX4 in MSCs could ameliorate adipose tissue and metabolic phenotypes during obesity. To this end, we generated MSC-selective *Sox4* knockout (*Sox4*-KO) mice by crossing *Pdgfra^CreERT2^* mice with *Sox4^fl/fl^* mice. Adult *Sox4*-KO mice and control littermates were treated with tamoxifen to delete *Sox4* (in KO mice) and subsequently fed a HFD for 8 weeks. Control and *Sox4*-KO mice exhibited similar body weights and comparable adipose tissue and liver weights (**Fig. 4a, b; Extended Data Fig. 5a**). Strikingly, *Sox4*-KO mice showed markedly improved glucose tolerance (**Fig. 4c**). This was accompanied by lower plasma insulin levels following glucose challenge (**Fig. 4d**).

**Fig. 4.**
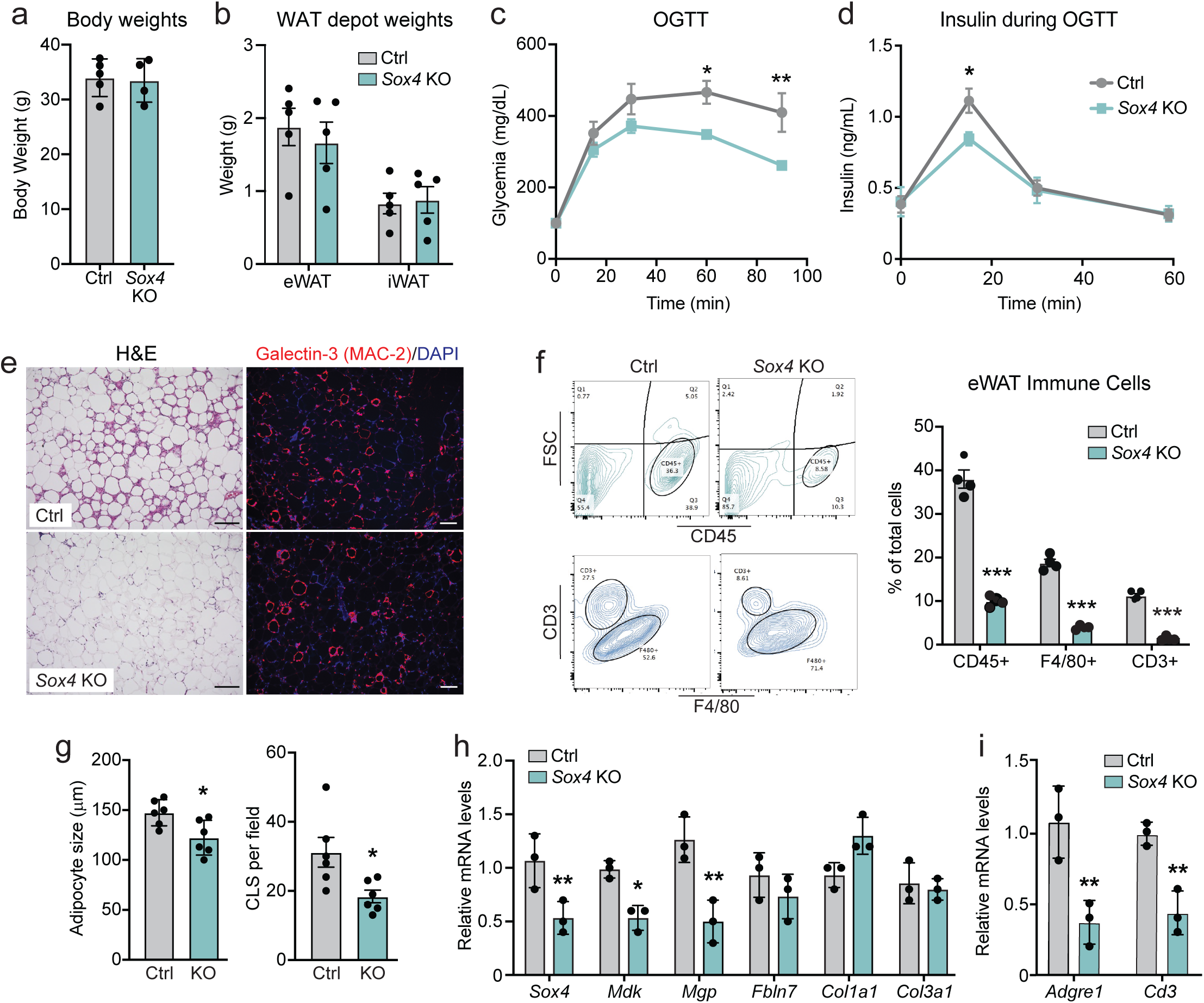
*Sox4* deletion in MSCs reduces adipose tissue inflammation and improves glucose homeostasis in obesity. *Pdgfra^CreERT2^*; *Sox4^LoxP^* (*Sox4*-KO) and control mice were treated with tamoxifen to delete *Sox4* in MSCs and then fed a HFD for 8 weeks. **(a)** Body weights. **(b)** iWAT and eWAT depot weights. **(c)** OGTT. **(d)** Plasma insulin levels during OGTT. **(e)** H&E staining and immunofluorescence staining of Galectin 3 (Mac2, red) and nuclei (DAPI, blue) in eWAT sections. **(f)** Quantification of adipocyte size and CLS per field in eWAT. **(g)** Flow cytometry gating strategy and quantification of CD45^+^ immune cells, ADGRE1 (F4/80)^+^ macrophages, and CD3^+^ T cells in eWAT. **(h)** mRNA levels of CAF-like cell genes (*Mdk*, *Mgp*, *Fbln7*, *Col1a1*, and *Col3a1*) in eWAT. **(i)** mRNA levels of *Adgre1* (macrophage) and *Cd3* (T cell) in eWAT (n = 4-5 mice per group). Data are presented as mean ± SEM. Statistical significance: *P < 0.05; **P < 0.01; ***P < 0.001.

Histological analysis showed that *Sox4*-KO mice had substantially improved eWAT morphology, with less inflammation and fibrosis and reduced adipocyte size (**Fig. 4e, g**). Immunostaining for the macrophage marker Galectin-3 (Mac-2) revealed significantly fewer crown-like structures in *Sox4*-KO eWAT, indicative of reduced macrophage infiltration (**Fig. 4e, g**). Flow cytometry analysis of stromal vascular cells revealed a marked reduction in total immune cells (CD45^+^), macrophages (F4/80^+^), and T cells (CD3^+^) in eWAT from *Sox4*-KO mice relative to controls (**Fig. 4f**). At the gene expression level, eWAT from *Sox4*-KO mice exhibited reduced expression of CAF-like cell genes (*Mdk*, *Mgp*, *Fbln7*, *Col1a1*, *Col3a1*), along with decreased expression of macrophage (*Adgre1*) and T cell (*Cd3e*) marker genes (**Fig. 4h, i**). No overt histological differences were observed in iWAT or liver from *Sox4*-KO mice relative to controls; however, expression of *Sox4* and CAF-associated, inflammatory, and matrix-remodeling genes were reduced or showed downward trends (**Extended Data Fig. 5b-d**). Together, these results indicate that *Sox4* ablation in MSCs attenuates pathological remodeling in eWAT and improves metabolic homeostasis.

### CAF-like cells signal via MDK to drive adipose inflammation and glucose intolerance

CAFs influence tumor growth through secreted signaling molecules. We sought to identify paracrine factors produced by CAF-like cells that regulate the adipose tissue microenvironment and remodeling. We applied the CellChat tool to identify candidate ligand-receptor interactions between MSCs and other cell types in adipose tissue from the scRNA-seq dataset. This analysis identified many signaling pathways that were enriched in MSCs under HFD conditions, including POSTN and IFNγ (IFN-II), which are known to be associated with fibrosis and inflammation (**Fig. 5a**). The most highly enriched pathway in MSCs under HFD was the growth factor Midkine (MDK) (**Fig. 5a**). MDK is predicted to interact with multiple receptors, including LRP1, NCL, SDC2, and ITGB5 (**Fig. 5b**; **Extended Data Fig. 6a, b**). *Mdk* mRNA expression was mainly restricted to CAF-like cells in eWAT and was strongly upregulated by HFD (**Fig. 5c; Extended Data Fig. 6c**).

**Fig. 5.**
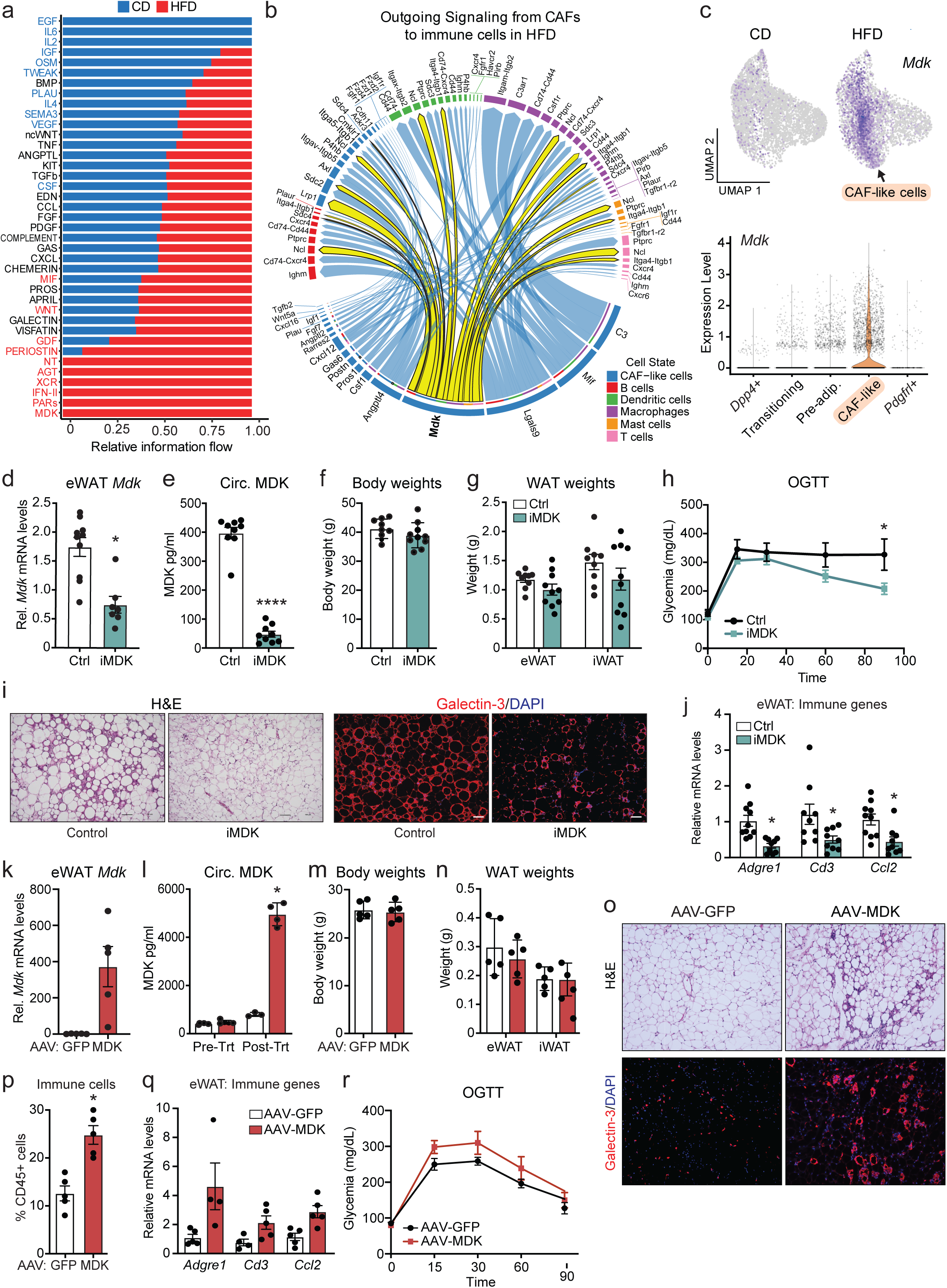
CAF-like cells signal via MDK to promote adipose tissue inflammation and impair glucose homeostasis. **(a**) CellChat analysis showing outgoing signaling pathways from CAF-like cells under CD or HFD conditions. **(b)** Predicted MDK ligand-receptor interaction network between CAF-like cells and immune cells under HFD conditions. **(c)** Feature plots and violin plot showing *Mdk* expression across MSC subsets. **(d-j)** C57BL/6 mice were fed a HFD for 16 weeks, followed by treatment with iMDK for 10 days (n = 9-10 mice per group). **(d)** *Mdk* mRNA levels in eWAT. **(e)** Circulating MDK levels. **(f)** Body weights. **(g)** eWAT and iWAT depot weights. **(h)** OGTT. **(i)** H&E staining and immunofluorescence staining of Galectin-3 (Mac2, red) and nuclei (DAPI, blue) in eWAT. **(j)** mRNA levels of inflammatory markers *Adgre1* (macrophage), *Cd3* (T Cell), and *Ccl2* in eWAT. **(k-r)** C57BL/6 mice maintained on CD received intra-eWAT injections of AAV-GFP or AAV-MDK (n = 5 mice per group). **(k)** *Mdk* mRNA levels in eWAT. **(l)** Circulating MDK levels. **(m)** Body weights. **(n)** eWAT and iWAT depot weights. **(o)** H&E staining and immunofluorescence staining of Galectin-3 (Mac2, red) and nuclei (DAPI, blue) in eWAT sections. **(p)** Flow cytometric quantification of CD45^+^ immune cells in eWAT. **(q)** mRNA levels of *Adgre1* (macrophage), *Cd3* (T cell), and *Ccl2* (inflammatory) in eWAT. **(r)** OGTT. All data are presented as mean ± SEM. Statistical significance: *P < 0.05.

To assess the function of MDK in obesity, we treated obese mice (fed a HFD for 16 weeks) with a commercially available MDK inhibitor (iMDK) for 10 days (**Extended Data Fig. 6d**). iMDK treatment decreased *Mdk* expression in eWAT as well as circulating MDK levels (**Fig. 5d, e**) without affecting body weight, adipose depot weights or liver weight (**Fig. 5f, g**; **Extended Data Fig. 6e**). Notably, MDK inhibition improved glucose tolerance (**Fig. 5h**). Histological analysis demonstrated that iMDK treatment markedly attenuated inflammation and fibrosis in eWAT (**Fig. 5i**). Consistent with these findings, qRT-PCR analysis of eWAT from iMDK-treated mice showed reduced expression of inflammatory and immune cell markers, including *Adgre1* (macrophages), *Cd3e* (T cells), and the chemokine *Ccl2*, compared with controls (**Fig. 5j**). iWAT from iMDK-treated mice showed reduced *Mdk* expression and a trend toward lower inflammatory gene expression without obvious tissue morphological changes (**Extended Data Fig. 6f, h**). In the liver, *Mdk* and inflammatory gene expression and histology were similar between iMDK-treated and control mice (**Extended Data Fig. 6g, i**). Together, these results suggest that beneficial remodeling of eWAT mediates the metabolic effects of MDK inhibition.

We also asked whether increased MDK expression is sufficient to promote pathological adipose tissue remodeling. To this end, we injected an AAV vector expressing either MDK or GFP (control) into the eWAT of lean mice (**Extended Data Fig. 6j**). AAV-MDK strongly induced *Mdk* expression in eWAT (**Fig. 5k**) and increased circulating MDK levels (**Fig. 5l**) without altering body weight or WAT depot weights (**Fig. 5m, n**). Histological analysis revealed inflammatory and fibrotic features in eWAT from mice receiving AAV-MDK (**Fig. 5o**). Consistent with these histological features, eWAT from AAV-MDK-treated mice contained more CD45^+^ immune cells than AAV-GFP controls (**Fig. 5p**) and expressed higher levels of the immune markers *Adgre1* (macrophages), *Cd3e* (T cells), and *Ccl2* (**Fig. 5q**). Additionally, even in the absence of metabolic challenge, AAV-MDK treatment produced a trend toward impaired glucose tolerance in mice maintained on a control diet (**Fig. 5r**). Together, these findings indicate that elevated MDK promotes inflammatory remodeling of eWAT and support a model in which MDK acts downstream of SOX4 to promote metabolic dysfunction.

### SOX4 and MDK are elevated in adipose tissue from individuals with T2D

To determine if *SOX4* and *MDK* may be associated with metabolic dysfunction in humans, we analyzed the expression of these genes in adipose tissue samples from BMI-matched obese individuals with or without T2D. Notably, both *SOX4* and *MDK* expression levels were significantly elevated in adipose tissue from T2D patients (**Fig. 6a**). In a publicly available human adipose tissue dataset, *SOX4* expression in both adipocytes and SVF positively correlated with BMI, whereas *MDK* showed little association in adipocytes but a strong positive trend in SVF ^27^ (**Extended Data Fig. 7a, b**).

**Fig. 6.**
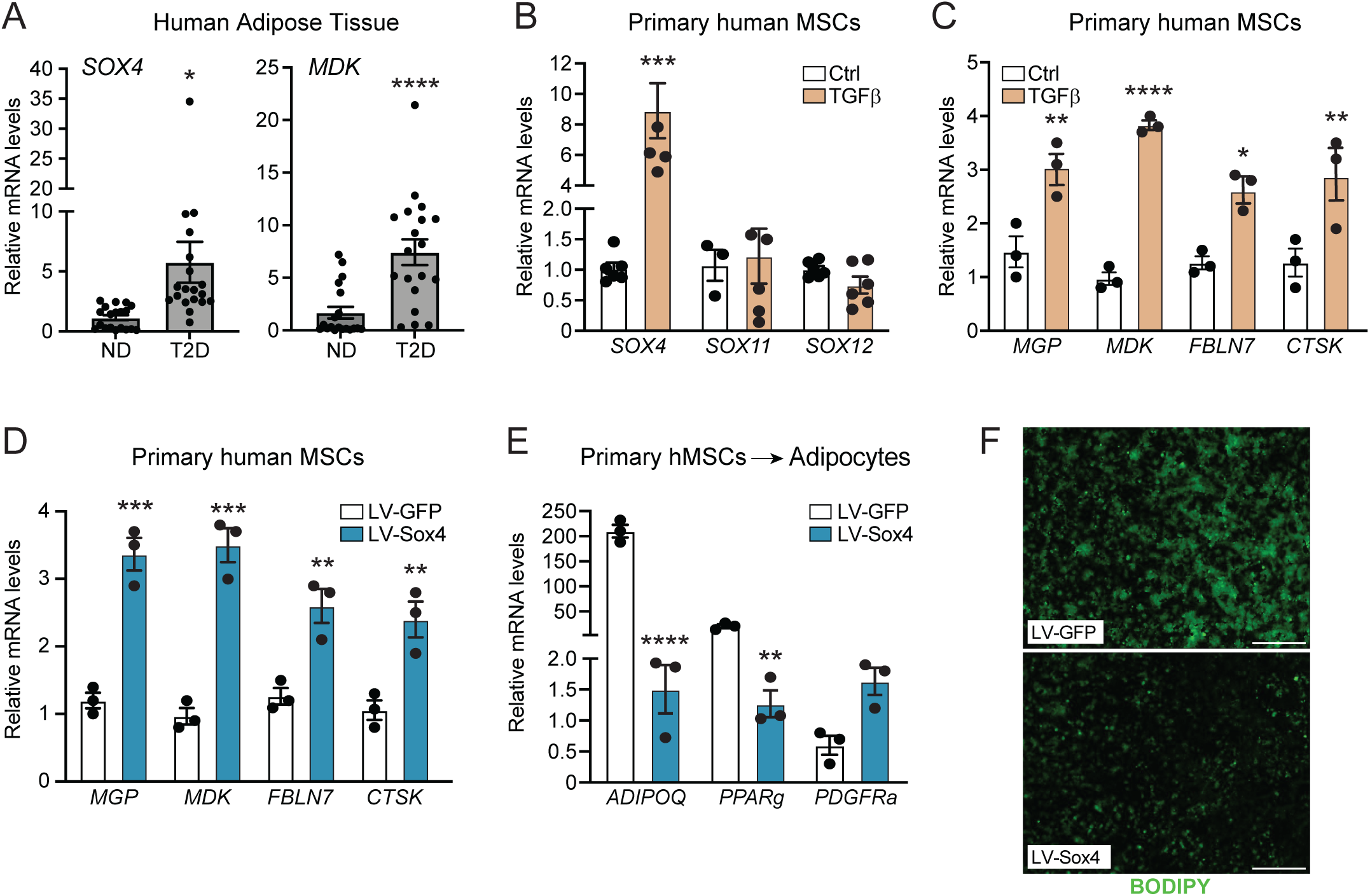
The SOX4-MDK CAF-like stromal program is conserved in human adipose cells. **(a)** mRNA levels *SOX4* and *MDK* in SVF of adipose tissue from BMI-matched obese individuals without diabetes (ND) and with type 2 diabetes (T2D) (n = 19 human donors per group). **(b)** mRNA levels of *SOX4*, *SOX11*, and *SOX12* in primary human MSCs treated with vehicle (Ctrl) or TGFβ (n = 5 per group). **(c)** mRNA levels of CAF-like cell genes *MGP*, *MDK*, *FBLN7*, and *CTSK* in primary human MSCs treated with vehicle (Ctrl) or TGFβ (n = 3 per treatment). **(d)** mRNA levels of CAF-like cell genes *MGP*, *MDK*, *FBLN7*, and *CTSK* in primary human MSCs transduced with lentiviral-GFP or SOX4 (n = 3 biological replicates per donor). **(e)** mRNA levels of adipocyte- selective genes *ADIPOQ* and *PPARG* and the MSC marker *PDGFRA* in GFP and SOX4-expressing MSCs (from (D)), induced to differentiate into adipocytes (n = 3 biological replicates per donor). **(f)** BODIPY staining of triglycerides in differentiated cultures from (E). All data are presented as mean ± SEM. Statistical significance: ns, not significant; *P < 0.05; **P < 0.01; ***P < 0.001; ****P < 0.0001.

To assess the potential relevance of the TGFβ-SOX4 pathway in humans, we treated primary MSCs isolated from subcutaneous adipose tissue with TGFβ, which robustly induced *SOX4* expression without affecting the related family members *SOX11* and *SOX12* (**Fig. 6b**). CUT&RUN analysis further revealed strong TGFβ-induced SMAD3 binding at the *SOX4* promoter, comparable to its recruitment to the canonical target gene *COL1A1*, whereas SMAD3 binding at the *SOX11* and *SOX12* loci was unchanged (**Extended Data Fig. 7c**). These results suggest that TGFβ signaling directly activates *SOX4* in human MSCs. TGFβ treatment of MSCs also upregulated CAF-associated genes, including *MDK*, *MGP*, *FBLN7*, and *CTSK* (**Fig. 6c**). Furthermore, lentiviral-mediated expression of SOX4 in human MSCs activated these CAF signature genes (**Fig. 6d**). SOX4 expression also impaired human adipocyte differentiation, as evidenced by reduced *ADIPOQ* and *PPARG* expression and diminished lipid accumulation following adipogenic induction (**Fig. 6e, f**). Together, these findings support a conserved role for a TGFβ-SOX4-MDK signaling axis in enforcing a non-adipogenic, CAF-like MSC phenotype that contributes to adipose tissue dysfunction in obesity and diabetes.

## Discussion

This study identifies MSCs as critical effectors of maladaptive adipose tissue remodeling in obesity, demonstrating that their reprogramming into a CAF-like state promotes inflammation and systemic metabolic dysfunction. Mechanistically, obesity-induced SOX4 activation redirects MSCs away from adipogenesis and toward a fibro-inflammatory phenotype. Modulating SOX4 activity in MSCs alters adipose tissue remodeling and systemic metabolism without affecting body weight or fat mass, demonstrating that MSC state can regulate metabolic health independently of adiposity.

Previous studies have identified fibro-inflammatory stromal populations in visceral adipose tissue, including LY6C⁺; PDGFRβ⁺ fibro-inflammatory progenitors described by Hepler et al. (2018) and CD9⁺ fibroblasts identified by Marcelin et al. (2017)^10,12^. Our results reveal that obesity additionally induces a molecularly distinct CAF-like state within the MSC compartment. These CAF-like cells are defined by a SOX4-driven transcriptional program enriched for inflammatory genes and by their marked expansion in obesity. Notably, Nahmgoong et al. identified a related stromal population in visceral WAT with an overlapping gene expression profile, providing independent support for the CAF-like state described here^28^.

Adipose tissue MSCs are heterogeneous and hierarchically organized, with DPP4⁺ cells residing upstream of more committed adipogenic cells^24–26^. Our data suggest that obesity redirects the output of this hierarchy. In this context, impaired adipogenesis may reflect not only defective differentiation, but active lineage reassignment toward a non-adipogenic, CAF-like fate. This is consistent with an emerging view that fibroblasts occupy conserved cross-tissue states and can adopt recurrent pathological programs in response to chronic stress or injury^26,29^.

We identified SOX4 as a central regulator of the MSC fate transition into CAF-like cells. SOX4 has been previously linked to cell plasticity and TGFβ-responsive mesenchymal programs in developmental and cancer contexts ^30–33^. SOX4-activation in MSCs impaired adipogenic competence while inducing CAF-associated gene expression, consistent with prior work showing that SOX4 suppresses pre-adipocyte determination and limits adipocyte hyperplasia ^34^. Beyond its anti-adipogenic effects, SOX4 drives a pro-inflammatory stromal niche in adipose tissue. Visceral eWAT, known to exhibit the most pronounced pathology in obesity, was most sensitive to SOX4 activity in MSCs, though modulation of SOX4 also altered CAF-like gene expression in subcutaneous iWAT and rpWAT. Thus, pathogenic MSC reprogramming may occur broadly across depots but require longer HFD exposure to become fully apparent.

An important aspect of this study was the identification of MDK as a paracrine effector of the CAF-like cells. Secretion of MDK enables a relatively restricted stromal population to exert broad effects on adipose tissue physiology. MDK is well positioned to mediate microenvironmental remodeling, as it has been implicated in inflammatory, fibrotic, and immune-regulatory responses across multiple pathological settings^35–38^. Consistent with a broader link between SOXC activity, MDK, and inflammatory stromal signaling, a related SOXC-dependent program has also been described in bone marrow MSCs. SOXC inactivation in *Lepr*⁺ bone marrow MSCs reduced expression of *Mdk* along with interferon and chemotaxis-associated genes^39^. These findings suggest that tissue-resident MSCs may adopt conserved SOXC-dependent inflammatory programs across distinct tissue contexts. Further supporting this model, MDK has been shown to activate CD8⁺ T cells, which promote myeloid cell recruitment and amplify inflammatory remodeling^40^. This MDK- T cell-myeloid signaling axis is particularly relevant to our findings, as SOX4 activation in MSCs increased T cell and macrophage markers in adipose tissue, whereas MDK inhibition had the opposite effect.

Importantly, *SOX4* and *MDK* expression were increased in adipose tissue stromal cells from people with type 2 diabetes. Circulating MDK levels have also been shown to positively correlate with BMI^41,42^. Furthermore, activation of TGFβ or SOX4 in human MSCs drives a CAF-like program and suppresses adipogenic activity, analogous to the effects observed in mice. Together, these findings suggest that the TGFβ-SOX4-MDK signaling axis in MSCs has translational relevance in humans.

A limitation of this study is the use of the *Pdgfra-CreERT2* model, which targets MSCs in multiple tissues, including adipose tissue. PDGFRα is a well-established marker of adipose MSCs, and *Pdgfra-CreERT2* currently represents one of the most effective and widely used genetic approaches to target this population in adult mice. Nevertheless, we cannot exclude contributions from extra-adipose MSCs to the systemic metabolic phenotypes observed in our SOX4 mouse models. The most pronounced effects of SOX4 manipulation, however, were observed in adipose tissue, particularly visceral eWAT, where changes in stromal cell identity were accompanied by marked alterations in inflammation and fibrosis. In contrast, no overt histological abnormalities were detected in the liver, despite moderate changes in inflammatory and fibrotic gene expression. Together, these findings support altered adipose tissue remodeling as a major contributor to the observed metabolic phenotypes, while highlighting the need for more genetic tools to define the specific contribution of adipose MSCs to systemic metabolic regulation.

In summary, this work identifies pathogenic MSC reprogramming as a driver of adipose tissue pathology and metabolic disease. Targeting stromal fate regulators or their paracrine outputs, such as SOX4 or MDK, may represent a therapeutic strategy to restore healthy adipose remodeling and ameliorate metabolic disease.

## Materials and Methods

### Mice

All animal experiments were performed in accordance with protocols approved by the University of Pennsylvania Institutional Animal Care and Use Committee (IACUC protocol no. 805649). Mice were maintained under standard housing conditions by the University of Pennsylvania University Laboratory Animal Resources, with a 12-h light/dark cycle and *ad libitum* access to water and either regular control diet (LabDiet, 5010) or high fat diet (HFD; Research Diets, D12492). Experiments were performed in male mice between 10 and 18 weeks of age unless otherwise indicated. For MSC- selective gene deletion or expression of SOX4, we used *Pdgfra^CreERT2^* mice from Jackson Laboratory (strain no. 018280). *Pdgfra^CreERT2^* mice were crossed with either *Sox4^LoxP^* mice^43^ or *Rosa26^Sox4^* mice^39^ to generate *Sox4*-KO and SOX4-OE mice, respectively. CRE recombinase activity was induced at 6 weeks of age by intraperitoneal administration of tamoxifen dissolved in corn oil (Sigma, T5648; 20 mg/mL stock solution) once daily for five consecutive days. For HFD studies, mice were placed on HFD one week after tamoxifen administration. *Sox4*-GFP reporter mice [B6;FVB-Tg(*Sox4*- EGFP)HW173Gsat/Mmucd] were purchased from MMRRC (strain no. 030033-UCD) and subjected to HFD for 12 weeks before analysis.

For pharmacological inhibition of MDK, mice received daily intraperitoneal injections of iMDK (MedChemExpress, CAS 881970-80-5) dissolved in corn oil at 9 mg/kg, or vehicle control, for 12 days. For MDK overexpression, AAV-MDK or AAV-GFP control virus was injected bilaterally into eWAT, and mice were analyzed 3 weeks after injection.

For oral glucose tolerance tests (OGTT), mice were fasted for 16 hours prior to oral administration of glucose (2g/kg body weight). Blood glucose was measured from tail vein blood at 0, 15, 30, 60 and 90 minutes after glucose administration using Ascensia Contour Next glucometer (Medline, USA). Tail blood was collected at similar time points for insulin levels measurements. Plasma insulin levels were measured using Crystal Chem 90080 ELISA kit according to the manufacturer’s instructions.

### Histology and Immunofluorescence

Tissues were fixed in 4% paraformaldehyde overnight, washed in PBS, dehydrated in ethanol, paraffin-embedded and sectioned. Slides were incubated in primary antibody overnight [(SOX4 (1:500, Invitrogen, MA5-31423); PDGFRa (1:500, R&D, AF1062); Galectin-3 (MAC2) (1:250, Invitrogen, 14-5301-82)] and secondary antibody conjugated to peroxidase and then developed using Tyramide Signal Amplification (TSA, Akoya Biosciences). Images were captured on Leica Stellaris or Keyence BZ-X700 fluorescent light microscope.

### Cell culture and adipogenic differentiation

#### Isolation of stromal vascular cells

WAT was dissected, minced gently and digested with Collagenase Type I (1.5 units/ml; Worthington) and Dispase II (2.4 units/ml; Roche) in DMEM/F12 containing 1% fatty acid-free bovine serum albumin (Gold Biotechnology) in a gentleMACS dissociator (Miltenyi Biotec) as described before^24^. The digestion was quenched with DMEM/F12 containing 10% FBS, and dissociated cells were passed through a 100 μm filter and spun at 400 x g for 4 minutes. The pellet was resuspended in red blood cell lysis buffer (BioLegend), incubated for 4 minutes at RT, then quenched with DMEM/F12 containing 10% FBS. Cells were passed through a 70 μm filter, spun, resuspended, then passed through a final 40 μm filter, spun at 400 x g for 4 minutes.

Primary stromal vascular cells and 3T3-L1 cells were maintained in DMEM supplemented with 10% FBS and 1% penicillin/streptomycin at 37°C and 5% CO₂. Adipogenic differentiation was induced by treating cells with adipogenic cocktail (500 μM IBMX, 10 μM dexamethasone, 125 μM indomethacin, 1 μM rosiglitazone (Caymen), and 20 nM insulin) for 2 days and then maintained for 6 additional days in maintenance medium (1 μM rosiglitazone, 20 nM insulin). In conditions without rosiglitazone, rosiglitazone was omitted from both induction and maintenance media. Where indicated, cells were infected with lentivirus expressing SOX4 or GFP for 72 h, with medium replaced 24 h after infection. For TGFβ stimulation, cells were treated with 5 ng/mL recombinant mouse TGFβ1 (R&D Systems) or vehicle control for 48 h. Adipogenesis was assessed by staining with Bodipy (Invitrogen catalog no. D3922) for lipid droplet accumulation and by staining with Hoechst (Thermo Fisher catalog no. 62249) for nucleus number at 4 to 6 days post-induction (mouse cells) and 14 to 18 days post-induction (human cells). The cultures were imaged on a Keyence inverted fluorescence microscope (BZX-710).

#### CRISPR-Cas9-mediated Sox4 targeting

3T3-L1 cells were engineered by nucleofection of Cas9-sgRNA ribonucleoprotein complexes using the Lonza 4D-Nucleofector system and the P1 Primary Cell 4D X Kit L (Lonza, V4XP-1024). Synthetic *Sox4* sgRNA was complexed with recombinant CAS9 protein and delivered using program CA-137. Following nucleofection, cells were transferred to complete growth medium, allowed to recover overnight, and cultured for an additional 2 days before downstream treatments.

### Viral vectors

The mouse *Mdk* coding sequence was cloned into an AAV vector using HiFi DNA assembly and verified by Sanger sequencing. AAV-MDK and AAV-GFP were produced in HEK293T cells by PEI-mediated transfection with the corresponding transfer plasmid, rep/cap plasmid, and adenoviral helper plasmid. Viral particles were harvested from culture supernatants 6 days after transfection, benzonase-treated, filtered, and concentrated by PEG8000/NaCl precipitation. AAV genome titers were determined by DNase-resistant qPCR. Lentiviral particles expressing MDK, SOX4, or GFP were generated by PEI-mediated transfection of HEK293T cells with the corresponding transfer vector, psPAX2 packaging plasmid, and pMD2.G envelope plasmid. Lentiviral supernatants were collected 48 and 72 h after transfection, filtered, concentrated using PEG-it precipitation.

### RNA Extraction, qRT-PCR and RNA Sequencing

Total RNA was extracted using Trizol (Invitrogen) combined with Purelink RNA columns (Fisher) and quantified using a Nano-drop. mRNA was reverse transcribed to cDNA using the ABI High-Capacity cDNA Synthesis kit (ABI). Real-time PCR was performed on a QuantStudio5 qPCR machine using SYBR green fluorescent dye (Applied Biosystems). Fold changes were calculated using the ddCT method, with 18s mRNA serving as a normalization control.

### Western Blot

Total cell lysates were prepared in RIPA buffer containing 150 mM NaCl, 1% NP-40, 0.1% sodium deoxycholate, 0.1% SDS, and 50 mM Tris-HCl, pH 7.4, supplemented with protease and phosphatase inhibitors (Roche). Lysates were sonicated, mixed with 4× NuPAGE LDS Sample Buffer containing 10% 2-mercaptoethanol, and denatured at 95°C for 5 min. Proteins were separated on NuPAGE Novex 4-12% Bis-Tris gels and transferred to nitrocellulose membranes. Membranes were incubated with HRP-conjugated anti-FLAG M2 antibody (mouse monoclonal, 1:1000; Millipore Sigma, A8592).

### Flow cytometry

DPP4⁺ and PDGFRα⁺ cells were isolated as previously described^24^. Briefly, stromal vascular cells from WAT were resuspended in FACS buffer (HBSS containing 3% FBS), then incubated for 1 h at 4°C with the following antibodies: CD26/DPP4-FITC (BioLegend, 137806; 1:200), anti-mouse CD45-APC/Cy7 (BioLegend, 103116; 1:1000), anti-mouse CD31-APC-Fire (BioLegend, 102528; 1:1000), F4/80-APC (BioLegend, 123116; 1:500), and CD3-Alexa Fluor 488, clone 17A2 (eBioscience, 53-0032-82; 1:400). DAPI (Roche, 10236276001; 1:10,000) was added during the final 5 min of staining. Cells were washed three times with FACS buffer. Cells were sorted using a BD FACSAria cell sorter equipped with a 100-μm nozzle. Compensation was performed at the time of acquisition in BD FACSDiva software using compensation beads (BioLegend, A10497) for single-color controls and unstained SVCs for negative and DAPI fluorescence controls. Final data were analyzed using FlowJo software.

### Human samples

Adipose tissues were collected from human donors as part of surgical procedures at the University of Pennsylvania. Male and female subjects over the age of 18 signed a written informed consent preoperatively. Tissue samples were collected intact and consisted of 1- to 10-kg intact blocks of discarded tissue, including skin, viscera and subcutaneous adipose. Tissues were processed within 1 to 3 hours of removal from the patient. Adipose tissue was dissected from the skin or viscera and immediately flash frozen. This work was performed under the approval of Institutional Review Board protocol 0063275 for The Human Metabolic Tissue Bank. For human correlation analyses, publicly available human adipose datasets from Emont et al. ^27^ were used to assess *SOX4* and *MDK* expression in MSCs as a function of BMI. Expression values were summarized at the sample level, plotted as log2(expression + 1), and analyzed using Pearson and Spearman correlations within each tissue compartment.

### Single-cell RNA sequencing and analysis

Stromal vascular cells were isolated from eWAT and loaded onto a 10x Genomics platform to generate single-cell barcoded libraries using the 10x Genomics 3′ v2 chemistry according to the manufacturer’s instructions. Libraries were sequenced on an Illumina HiSeq 2500 platform, and raw sequencing data were processed with Cell Ranger v4.0.0 using the mm10 reference genome. Raw count matrices were processed in R using Seurat. Empty droplets were removed using DropletUtils, and ambient RNA contamination was corrected with SoupX. Cells with low gene complexity, high UMI counts, or high mitochondrial content were excluded. After quality control, data were log-normalized, variable genes were identified, and samples were integrated using Seurat canonical correlation analysis. Cell-cycle scores and mitochondrial transcript percentage were regressed during scaling. Dimensionality reduction was performed by PCA, neighbor graph construction, clustering, and UMAP visualization. Clusters were annotated using canonical marker genes, and differentially expressed genes were identified using Seurat’s Wilcoxon rank-sum test. MSCs (marked by: *Pdgfra*, *Col3a1*, *Dpt*, *Serping1*, *Plpp3*, and *Col1a2*) were re-clustered using the integrated assay. MSC subclusters were annotated based on marker gene expression.

Pseudotime trajectories were inferred using Slingshot, with the *Dpp4*^+^ progenitor cluster set as the root. Genes associated with pseudotime were identified using generalized additive modeling, and dynamic expression patterns were visualized as z-score-normalized heatmaps. RNA velocity was estimated using velocyto and scVelo, and trajectories were visualized on UMAP embeddings. Cell- cell communication analysis was performed using CellChat with the mouse ligand-receptor database restricted to secreted signaling pathways.

### Quantification and statistical analysis

Data are presented as mean ± SEM unless otherwise indicated. Statistical significance was determined using unpaired two-tailed Student’s t-test or ANOVA with appropriate post hoc tests. A p value < 0.05 was considered statistically significant.

### Data, code, and materials availability

Sequencing data have been deposited at GEO under accession number GSE333670. The data was processed using published software tools, for which detailed parameters are described in the materials and methods section.

## Acknowledgments

We are grateful to the single cell technology core (Children’s Hospital of Philadelphia Research Institute). We thank Jeff Ishibashi (UPenn) for valuable input on the project. We are grateful for funding from: National Institutes of Health grant DK120982 (PS), National Institutes of Health grant DK019525 (Penn Diabetes Center), Rubicon grant 40-45200-98-21118 from the Netherlands Organization for Health Research and Development (JMET), American Diabetes Association postdoctoral fellowship (KD, LL).

## Author contributions

KD and PS were responsible for project conceptualization and overall study design. KD and RPC performed the majority of the experiments and analyzed the results. CEMP, LC and LL performed experiments. KD and RPC generated visualizations. DMM, ST, MA, JMET and VL provided key reagents, samples and methodological expertise. KD and PS managed and supervised the project. PS acquired funding for the study. KD, PS and RPC wrote the manuscript.

## Competing interests

Authors declare that they have no competing interests.

## Notes

### Competing Interest Statement

The authors have declared no competing interest.

