## Extended Figures for "SOX4 Reprograms Adipose Stromal Cells into a Cancer-Associated Fibroblast- like State to Drive Metabolic Disease"

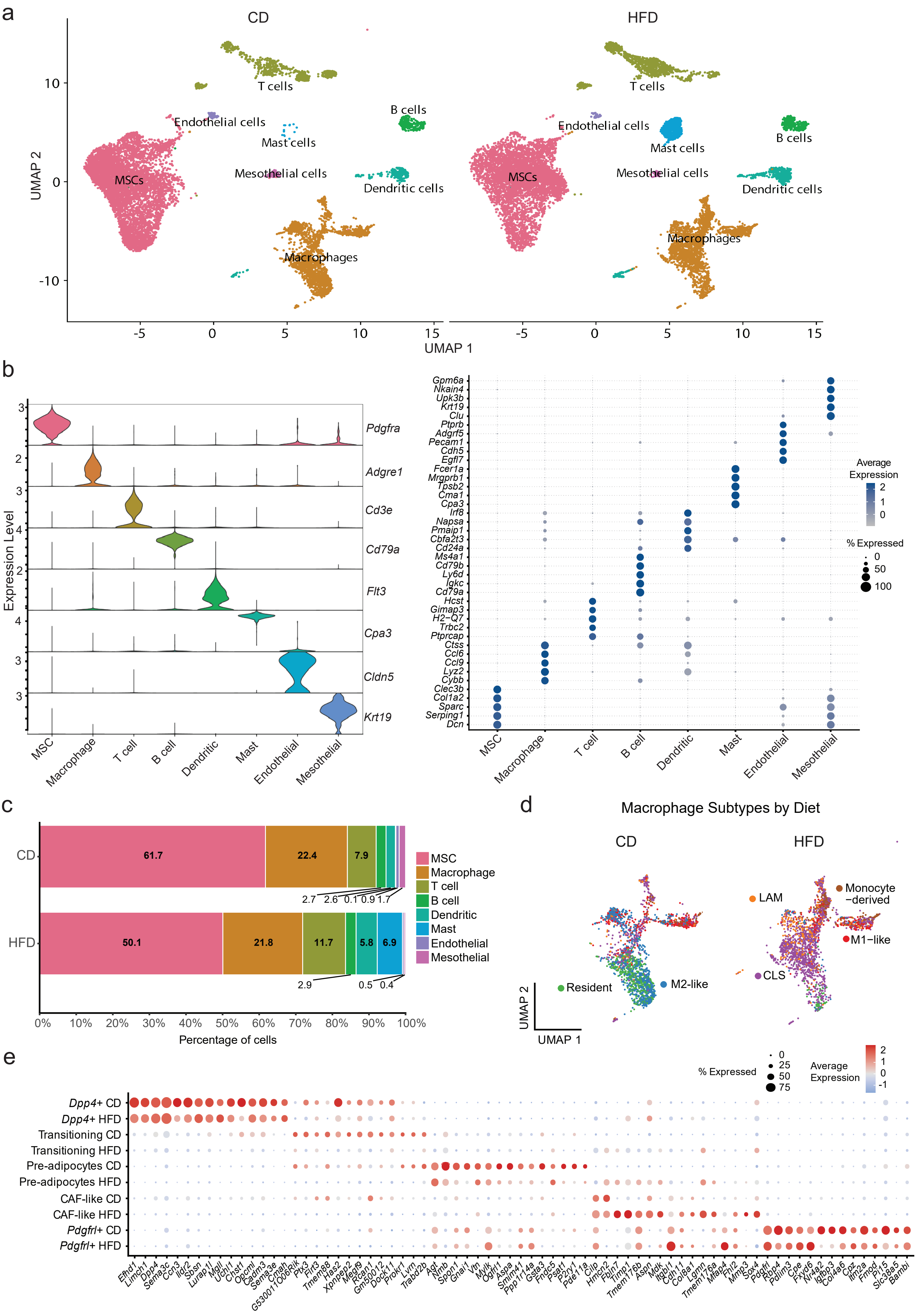
a
CD
HFD
T cells
B cells
Endothelial cells
Mast cells
Mesothelial cells
Dendritic cells
MSCs
Macrophages
UMAP 2
UMAP 1
10
0
-10
-5
5
15
b
Expression Level
Pdgfra
Adgre1
Cd3e
Cd79a
Flt3
Cpa3
Cldn5
Krt19
MSC
Macrophage
T cell
B cell
Dendritic
Mast
Endothelial
Mesothelial
Gpm6a
Nkain4
Upk3b
Krt19
Clu
Ptprb
Adgrf5
Pecam1
Cdh5
Egfl7
Fcer1a
Mrgprb1
Tpsb2
Cma1
Cpa3
Irf8
Napsa
Pmaip1
Cbfa2t3
Cd24a
Ms4a1
Cd79b
Ly6d
Igkc
Cd79a
Hcst
Gimap3
H2−Q7
Trbc2
Ptprcap
Ctss
Ccl6
Ccl9
Lyz2
Cybb
Clec3b
Col1a2
Sparc
Serping1
Dcn
Average Expression
% Expressed
c
CD
HFD
61.7
22.4
7.9
2.7
2.6
0.1
0.9
1.7
50.1
21.8
11.7
5.8
6.9
2.9
0.5
0.4
MSC
Macrophage
T cell
B cell
Dendritic
Mast
Endothelial
Mesothelial
0%
10%
20%
30%
40%
50%
60%
70%
80%
90%
100%
Percentage of cells
d
Macrophage Subtypes by Diet
CD
HFD
Resident
M2-like
LAM
Monocyte −derived
M1−like
CLS
UMAP 2
UMAP 1
e
% Expressed
Average Expression
Dpp4+ CD
Dpp4+ HFD
Transitioning CD
Transitioning HFD
Pre-adipocytes CD
Pre-adipocytes HFD
CAF-like CD
CAF-like HFD
Pdgfrl+ CD
Pdgfrl+ HFD
Efhd1
Limch1
Dpp4
Sema3c
Ccn3
Ildr2
Sbsn
Lurap1l
Mgll
Uchl1
Chst1
Opcml
Cadm3
Sema3e
Cnah
G530011O06Rik
Pi16
Flrt3
Tmem88
Has2
Xpnpep2
Megf9
Rcan1
Gm50012
Dock11
Prokr1
Lum
Trabd2b
Agt
Nmb
Spon1
Gnai1
Vtn
Mylk
Ogfrl1
Aspa
Smim41
Ppp1r14a
Gsta3
Fndc5
Psat1
P2ry1
Pde11a
Clip
Hmcn2
Fbln7
Timp1
Tmem176b
Aspn
Mdk
Itgbl1
Cdh11
Col8a1
Lgmn
Tmem176a
Mfap4
Fhl2
Mmp3
Sox4
Pdgfrl
Rbp4
Pdlim3
Cpe
Fxyd6
Nr4a2
Igfbp3
Col4a6
Cpz
Itm2a
Fmod
Pi15
Slc38a5
Bambi

f

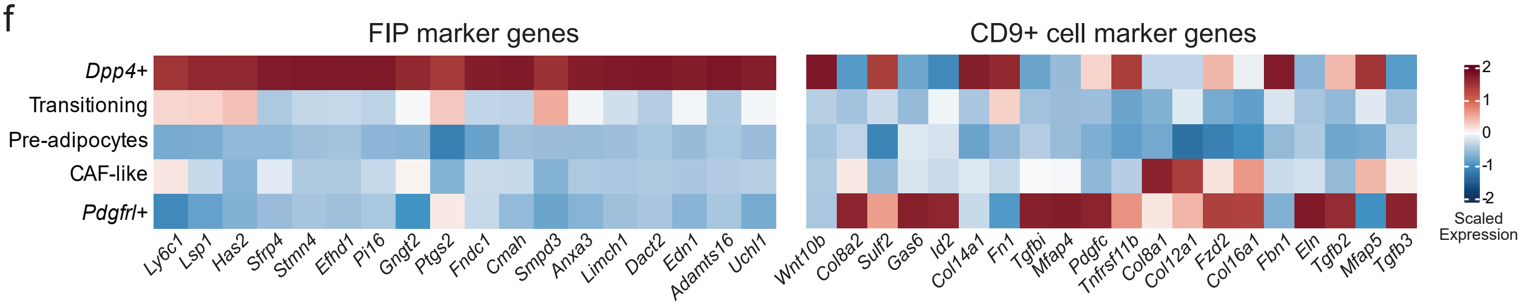


**Extended Data Fig. 1. Obesity remodels the cellular composition of adipose tissue and induces CAF-like cell genes across fibroblast states (related to Fig. 1).**

**(a)** UMAP representation of SVF cells from epididymal white adipose tissue (eWAT) of mice fed CD or HFD, split by diet condition and annotated by major cell types. **(b)** Violin plot and dot plot showing expression of marker genes used to annotate major SVF populations. **(c)** Stacked bar plot showing the relative abundance of major SVF cell populations under CD and HFD conditions. **(d)** UMAP of macrophage subclusters split by diet, identifying resident macrophages, M2-like macrophages, M1-like macrophages, LAMs, CLS-associated macrophages, and monocyte-derived macrophages. **(e)** Dot plot showing expression of cluster-enriched genes across MSC subsets split by diet. n = 3 mice per group pooled for scRNA-seq analysis. Dot size indicates the percentage of cells expressing each gene, and color indicates scaled average expression. Abbreviations: CD, CLS, crown-like structure; chow diet; eWAT, epididymal white adipose tissue; HFD, high-fat diet; LAM, lipid-associated macrophages; SVF, stromal vascular fraction. **(f)** Heatmap showing scaled expression of top marker genes for previously described fibro-inflammatory progenitors (FIPs) and CD9$^{+}$ fibroblasts across *Dpp4*$^{+}$ MSCs, transitioning state cells, pre-adipocytes, CAF-like cells, and *Pdgfrl*$^{+}$ fibroblasts.

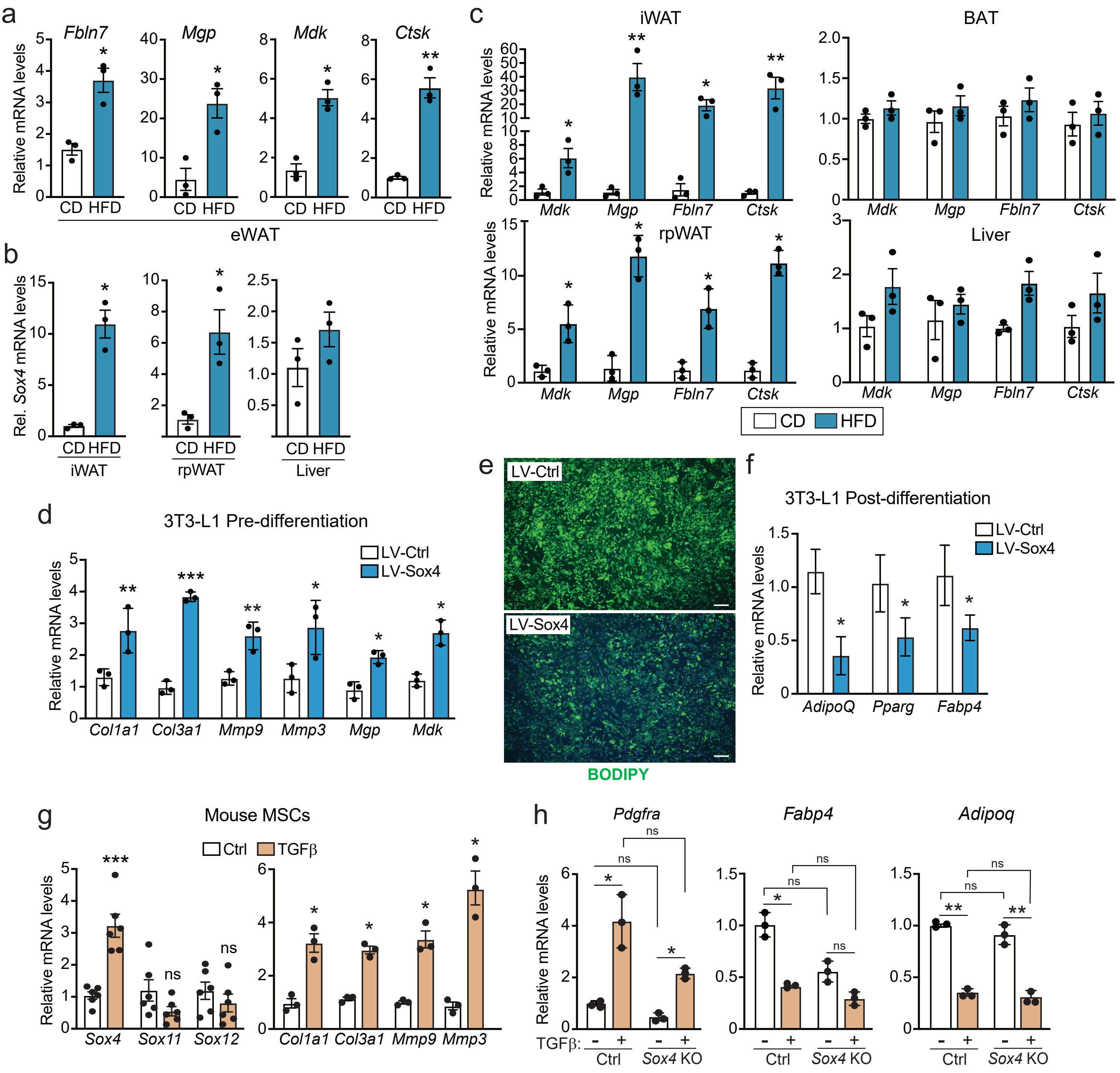


**Extended Data Fig. 2. HFD and TGFβ induce a SOX4-associated CAF-like gene program in adipose MSCs (related to Fig. 2)**

**(a-c)** C57BL/6 mice were fed CD or HFD for 16 weeks (n = 3 mice per group). **(a)** mRNA levels of CAF-like cell marker genes *Fbln7*, *Mgp*, and *Mdk* in eWAT. **(b)** mRNA levels of *Sox4* in iWAT, rpWAT and liver. **(c)** mRNA levels of CAF-like cell marker genes in iWAT, rpWAT, BAT, and liver. **(d)** mRNA levels of extracellular matrix and CAF-like cell marker genes in 3T3-L1 cells transduced with lentivirus (LV) expressing GFP (Ctrl) or SOX4. (n = 3 per condition). **(e)** BODIPY staining of triglycerides in Ctrl- and SOX4-expressing 3T3-L1 cells induced to undergo adipocyte differentiation. **(f)** mRNA levels of adipocyte marker genes in differentiated cultures from (E) (n = 3 per condition). **(g)** mRNA levels of *Sox4*, *Sox11*, *Sox12*, and fibrosis-related genes in mouse MSCs treated with TGFβ or vehicle control (ctrl) (n = 3-6 isolates per group). **(h)** mRNA levels of MSC marker *Pdgfra*, and adipocyte markers (*Fabp4* and *Adipoq*) in *Sox4*-KO and control (Ctrl) cells treated with TGFβ or vehicle (n = 3 per condition). All data are presented as mean +/- SEM. Statistical significance was determined by two-tailed unpaired Student's t test for two-group comparisons or two-way ANOVA for multi-condition comparisons. Statistical significance: ns, not significant; *P < 0.05; **P < 0.01; ***P < 0.001. Abbreviations: BAT, brown adipose tissue; CD, control diet; eWAT, epididymal white adipose tissue; HFD, high-fat diet; iWAT, inguinal white adipose tissue; qRT-PCR; rpWAT, retroperitoneal white adipose tissue.

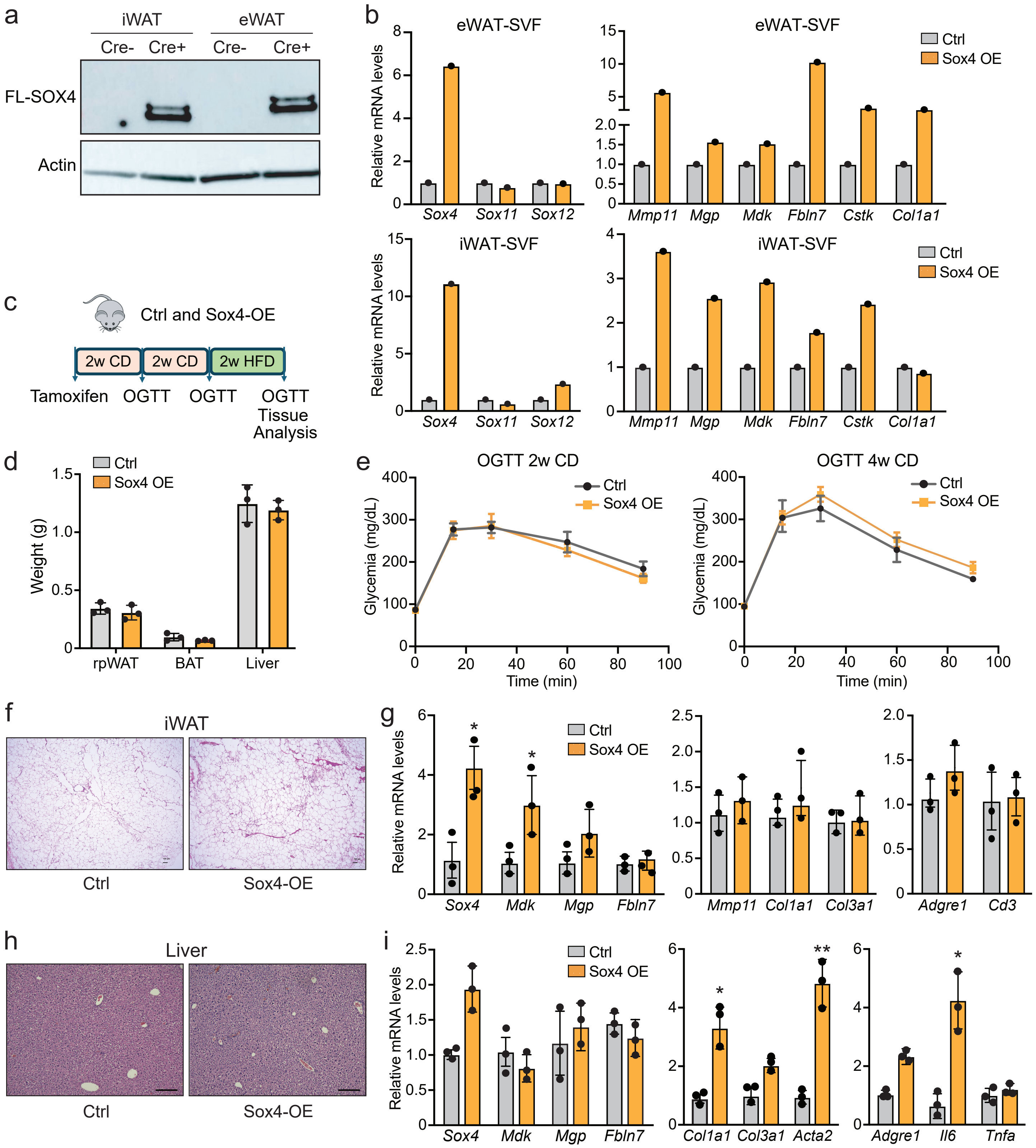


**Extended Data Fig. 3. Validation and tissue-level characterization of MSC-SOX4 activation (related to Fig. 3)**

**(a)** Immunoblot analysis of FLAG-SOX4 protein levels in isolated SVF cells from eWAT and iWAT of SOX4-OE and control mice following tamoxifen induction. Actin was used as a loading control. **(b)** mRNA levels of *Sox4*, *Sox11*, *Sox12*, and CAF-like cell marker genes in SVF cells from eWAT and iWAT of control and SOX4-OE mice (n = 2 independent plated SVF pools per group). **(c)** Schematic of experimental timeline for analyses of SOX4-OE and control mice in (D-I) (n = 3 mice per group). **(d)** Weights of rpWAT, BAT, and liver at endpoint. **(e)** OGTT after 2 and 4 weeks of CD feeding. **(f)** H&E staining of iWAT. **(g)** mRNA levels of *Sox4*, other CAF-like cell genes, and inflammatory marker genes in iWAT. **(h)** H&E staining of liver. **(I)** mRNA levels of *Sox4*, CAF-like cell genes, and inflammatory genes in liver. All data are presented as mean +/- SEM. Statistical significance was determined by two-tailed unpaired Student's t test or two-way ANOVA. Statistical significance: *P < 0.05; **P < 0.01. Abbreviations: BAT, brown adipose tissue; CD, control diet; eWAT, epididymal white adipose tissue; HFD, high-fat diet; iWAT, inguinal white adipose tissue; OGTT, oral glucose tolerance test; rpWAT, retroperitoneal white adipose tissue; SVF, stromal vascular fraction.

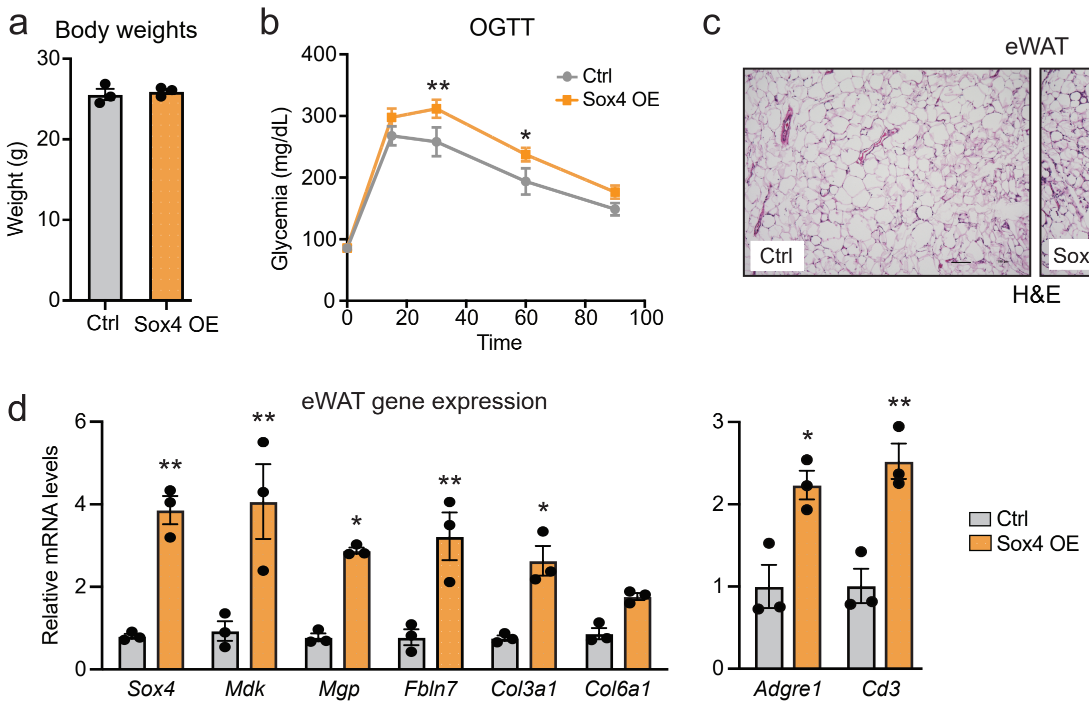


**Extended Data Fig. 4. MSC-SOX4 activation promotes eWAT inflammation and impairs glucose homeostasis (related to Fig. 3)**

*Pdgfra*$^{CreERT2}$; *Rosa26*$^{Sox4}$ (Sox4-OE) and control mice were treated with tamoxifen to activate SOX4 expression in MSCs, maintained on CD for 4 weeks and switched onto HFD for 2 weeks. **(a)** Body weights. **(b)** Oral glucose tolerance test. **(c)** H&E staining of eWAT. **(d)** mRNA levels of *Sox4*, CAF-like cell genes, and inflammatory genes in eWAT. Data are presented as mean +/- SEM. Statistical significance was determined by two-tailed unpaired Student's t test or two-way ANOVA. Statistical significance: *P < 0.05; **P < 0.01. Abbreviations: OGTT, oral glucose tolerance test.

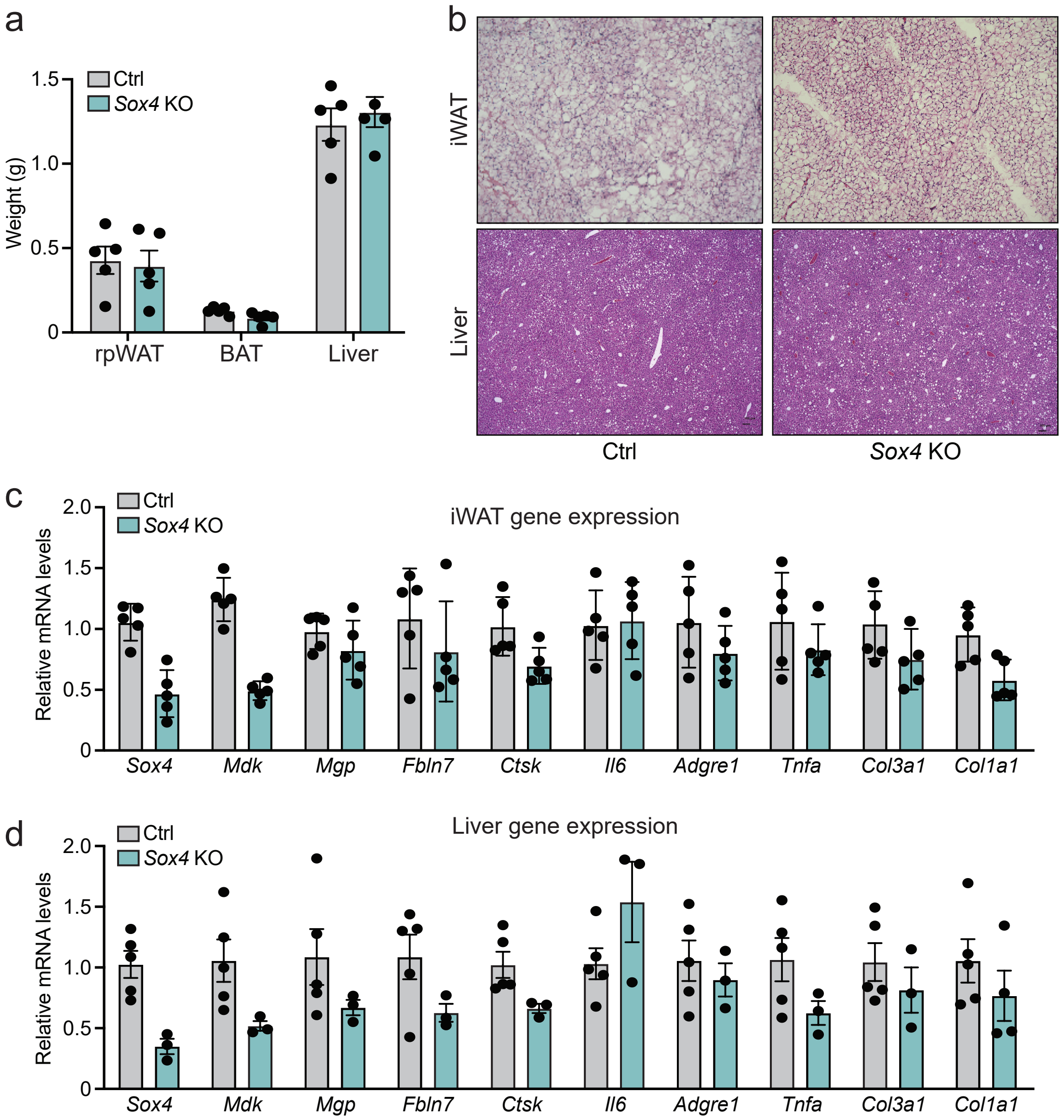


**Extended Data Fig. 5. Sox4 deficiency in MSCs does not markedly alter iWAT or liver morphology (related to Fig. 4)**

MSC-selective *Sox4*-KO mice and control mice were treated with tamoxifen and fed a HFD for 8 weeks (n = 4/5 mice per group). **(a)** Weights of rpWAT, BAT, and liver. **(b)** H&E staining of iWAT and liver sections. **(c,d)** mRNA levels of *Sox4*, other CAF-like cell and fibrotic genes, and inflammatory genes in iWAT (c) and liver (d). All data are presented as mean +/- SEM. Statistical significance was determined by two-tailed unpaired Student's t test. Abbreviations: BAT, brown adipose tissue; HFD, high-fat diet; iWAT, inguinal white adipose tissue; rpWAT, retroperitoneal white adipose tissue.

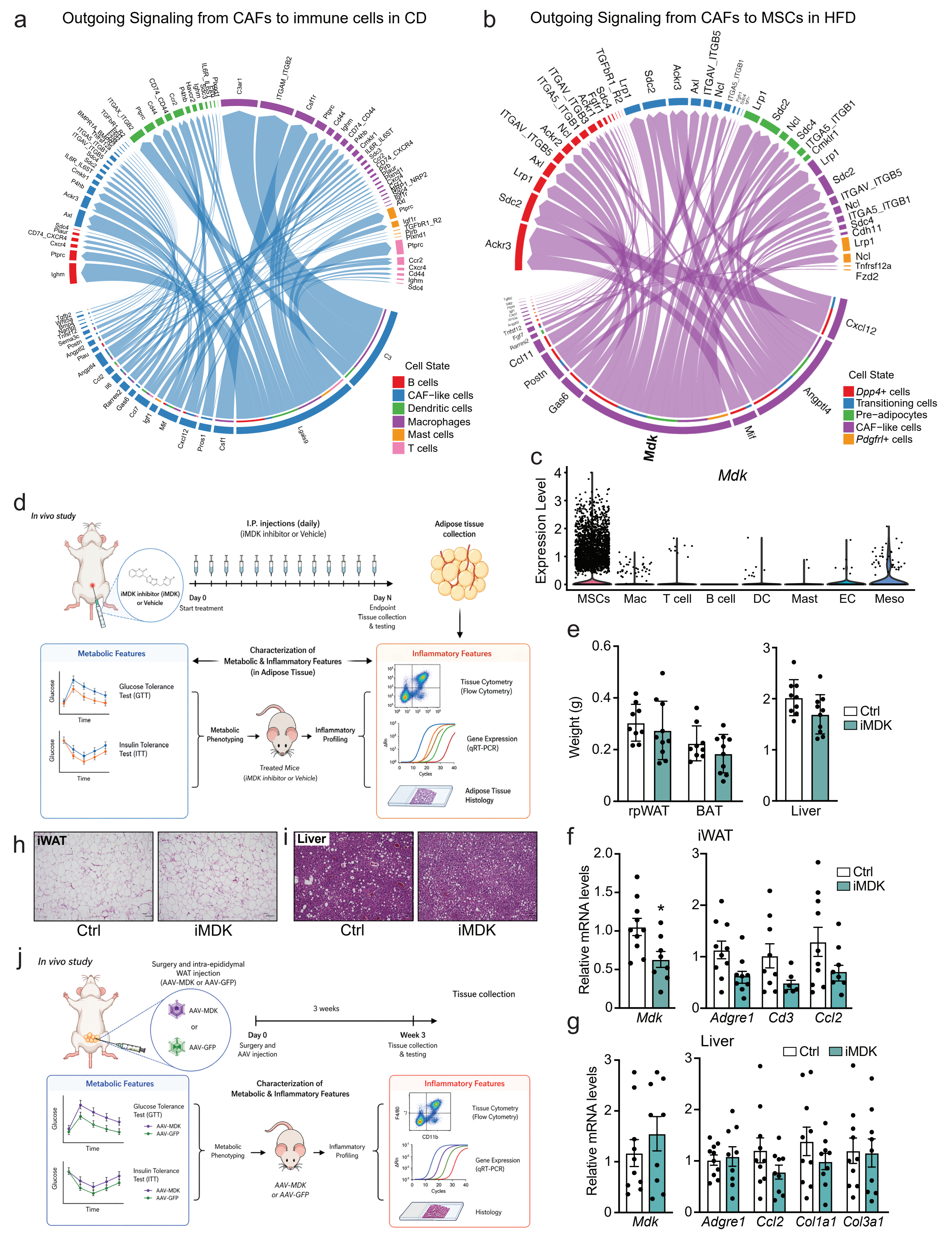


**Extended Data Fig. 6. CAF-like cell communication network and strategies to perturb MDK signaling (related to Fig. 5).**

**(a,b)** CellChat chord diagram s showing predicted ligand -receptor interactions from CAF -like cells to: immune cells under CD conditions (a) and MSCs under HFD conditions (b). **(c)** Violin plot showing *Mdk* expression across major SVF cell populations. **(d-i)** Obese mice (16 weeks of HFD) were treated with iMDK or vehicle (ctrl) for 10 days (n = 9-12 mice per group). **(d)** Schematic of experiments. **(e)** Weights of rpWAT, BAT, and liver **(f, g)** mRNA levels of *Mdk* and inflammatory genes in iWAT (f) and liver (g). **(h, i)** H&E staining of iWAT (h) and liver (i). **(j)** Schematic of AAV experiments. All data are presented as mean +/ - SEM. Statistical significance was determined by two-tailed unpaired Student's t test. Statistical significance: *P < 0.05. Abbreviations: AAV, adeno-associated virus; iMDK, MDK inhibitor; mac, macrophage; dc, dendritic cell; ec, endothelial cell; meso, mesothelial cell.

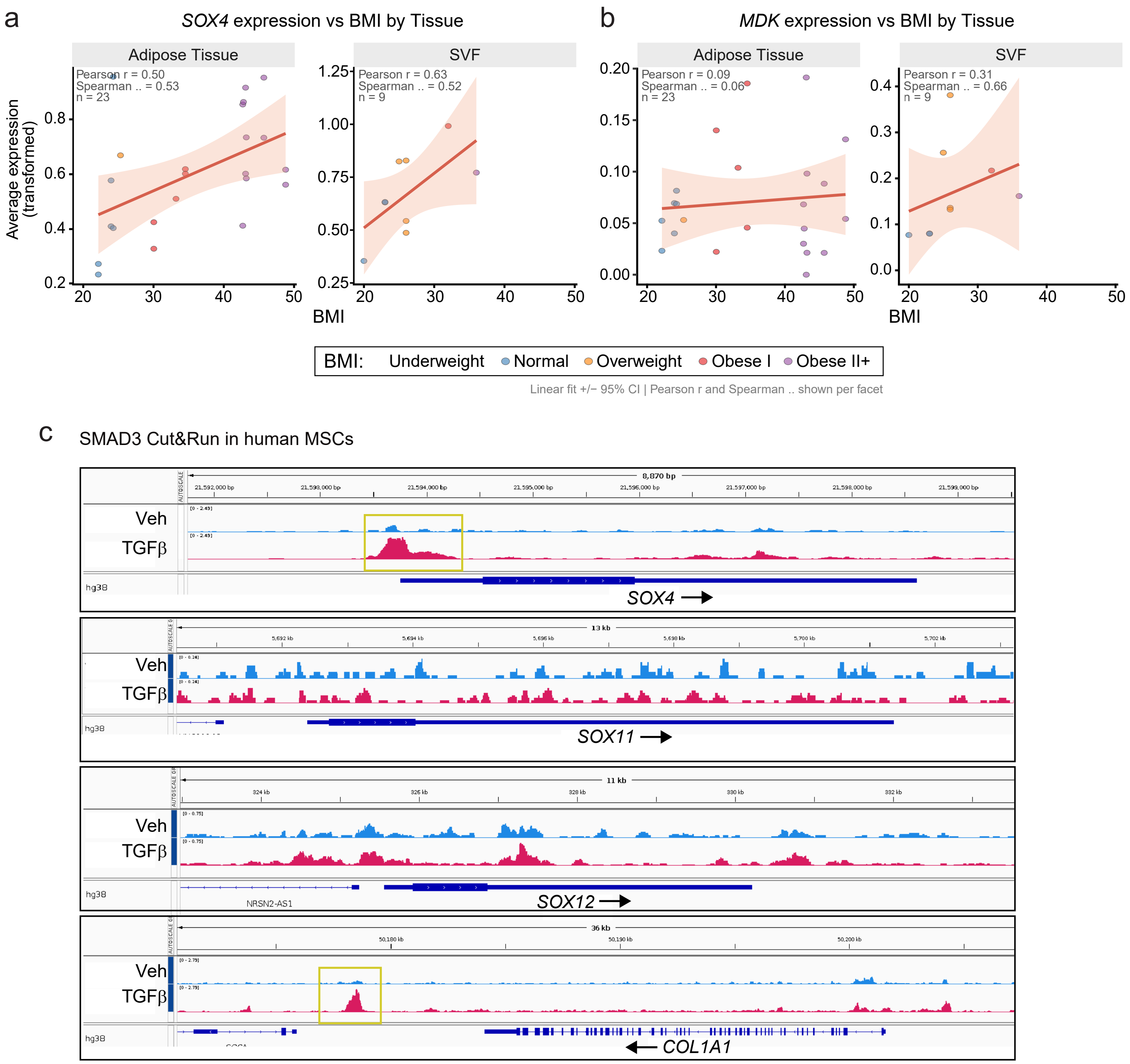


**Extended Data Fig. 7. *SOX4* and *MDK* expression positively correlate with BMI in human adipose tissue (related to Fig. 6).**

**(a,b)** Correlation analysis of *SOX4* (a) and *MDK* (b) expression with BMI in human adipose tissue and SVF cells. Linear regression is shown with 95% confidence intervals. Pearson and Spearman correlation coefficients are indicated for each tissue compartment (n = 23 adipose samples and n = 9 SVF samples). Samples are colored by BMI category. **(c)** Genome browser tracks showing SMAD3 binding at the indicated loci in primary human MSCs isolated from subcutaneous WAT. Abbreviations: BMI, body mass index; SVF, stromal vascular fraction.
